# Coupled aging of cyto- and myeloarchitectonic atlas-informed gray and white matter structural properties

**DOI:** 10.1101/2024.11.25.625295

**Authors:** David A. Hoagey, Ekarin E. Pongpipat, Karen M. Rodrigue, Kristen M. Kennedy

## Abstract

A key aspect of brain aging that remains poorly understood is its high regional heterogeneity and heterochronicity. A better understanding of how the structural organization of the brain shapes aging trajectories is needed. Neuroimaging tissue “types” are often collected and analyzed as separate acquisitions, an approach that cannot provide a holistic view of age-related change of the related portions of the neurons (cell bodies and axons). Because neuroimaging can only assess indirect features at the gross macrostructural level, incorporating post-mortem histological information may aid in better understanding of structural aging gradients. Longitudinal design, coupling of gray and white matter (GM, WM) properties, and a biologically informed approach to organizing neural properties are needed. Thus, we tested aging of the regional coupling between GM (cortical thickness, surface area, volume) and WM (fractional anisotropy, mean, axial, and radial diffusivities) structural metrics using linear mixed effects modeling in 102 healthy adults aged 20-94 years old, scanned on two occasions over a four-year period. The association between age-related within-person change in GM morphometry and the diffusion properties of the directly neighboring portion of white matter were assessed, capturing both aspects of neuronal health in one model. Additionally, we parcellated the brain utilizing the histological-staining informed von Economo-Koskinas atlas to consider regional cyto- and myelo-architecture. Results demonstrate several gradients of coupled association in the age-related decline of neighboring white and gray matter. Most notably, gradients of coupling along the heteromodal association to sensory axis were found for several areas (e.g., anterior frontal and lateral temporal cortices, vs pre- and post-central gyrus, occipital, and limbic areas), in line with heterochronicity and retrogenesis theories of aging. Further effort to bridge across data and measurement scales will enhance understanding of the mechanisms of the aging brain.

## 1 INTRODUCTION

Current understanding of development and aging of brain structure stems from both *in vivo* neuroimaging research over the last four decades and *ex vivo* histological research over the last 120 years. Particularly, structural MRI techniques have enabled and accelerated our understanding that aging is characterized by vast individual differences and regional heterogeneity across the brain’s macroscale, that underpin cognitive abilities. Histological techniques have historically revealed differences in microstructural organization in gray and white matter tissue, including the underlying neuronal composition, organization, and layering patterns. A deeper understanding of the biological, mechanistic factors underpinning age-related differences and changes to the brain’s structure could be gained from studies more frequently combining knowledge gleaned from *in vivo* macroscale MRI and *ex vivo* microscopy techniques. Further, the nature of imaging techniques exaggerates the separation of the neuronal parts into distinct tissue compartments. The current study aimed to add knowledge to this gap by leveraging pre-established, histologically derived cytoarchitectonic and myeloarchitectonic properties to inform aging of macroscale gray and white matter neuroimaging metrics, and to begin to “re-couple” the neuron’s parts, rather than treating them as separate gray and white matter compartments.

Brain aging is regionally heterochronic, with patterns of development and degradation, differing both across individuals and within-individuals over time. Several theoretical frameworks of structural brain aging have emerged, each describing differential susceptibility of brain regions and properties to the aging process, including prefrontal vulnerability (Dempster, 1992; Raz et al., 1997; West, 1996) anterior to posterior gradient (Bennett et al., 2010; Davis et al., 2009; Head et al., 2004; Madden et al., 2008; Pfefferbaum et al., 2005; Salat et al., 2005), and superior to inferior gradient (Hoagey et al., 2019; Sexton et al., 2014). More recent, integrated evidence points to a generalized vulnerability of association cortices, including the frontal but also the parietal (Fjell et al., 2009; Grieve et al., 2005; Sowell et al., 2003), and select temporal (Bartzokis et al., 2001; Fjell et al., 2009; Sowell et al., 2003) regions, as compared to visual, sensorimotor, or limbic cortices (Grieve et al., 2005; Salat et al., 2005; Ziegler et al., 2010). This observed association-sensory pattern of aging is often described as having a retrogenesis-like time course of differential vulnerability, such that the earliest regions to mature, from either a developmental or evolutionary perspective, are the latest regions to decline in aging (Raz, 2000; Salat et al., 2004). Taking this aging pattern into consideration with recent developmental literature characterizing cortical maturation in sensory-motor and associative regions (Sydnor et al., 2021), we believe there is also support for an aging sensorimotor-association axis that contrasts unimodal, inflexible, and perceptive regions with multi-modal, plasticity-derived, and cognitive center regions.

These differential patterns of age-related cortical decline associated with regional heterochronicity likely reflect differences in the underlying biological tissue properties (Koenig et al., 2022; Raz & Daugherty, 2017), which are difficult to assess with typical *in-vivo* MRI sequences (i.e., due to restrictions in scanning duration and sequence resolution). As such, these more fine-grained features of brain biology driving patterns of regional differentiation are poorly understood. Examination of laminar features, such as cytoarchitectural and myeloarchitectural patterns of organization, typically requires microscopy techniques in *ex-vivo* samples. Fortunately, microscopy-based brain research delineating regional differentiation in brain structure has been a major focus of neuroscientists since pioneering researchers began characterizing and parcellating the brain in the late 19^th^ and early 20^th^ centuries. Using *ex-vivo* dissection, combined with histological staining and macro-photography techniques, neuroanatomists have revealed microscopic tissue properties of cell size, laminar thickness, and neuronal organization, in an attempt to define areal boundaries (Brodmann, 1909), chronicle myelin precedence (Flechsig, 1920), or map regional cortical layering patterns (von Economo & Koskinas, 1925). These seminal efforts have resulted in several structural parcellation atlases that are still widely used today to differentiate and classify anatomical features based upon cytoarchitectonic and myeloarchitectonic patterns as well as provide a spatial and conceptual framework to map findings across studies. Recent work has pushed to digitize parcellation atlases to allow for direct mapping of the microstructural organization of the cortex with neuroimaging data (Markello et al., 2022; Pijnenburg et al., 2021; Scholtens et al., 2018). The von Economo - Koskinas (vE-K) atlas (Triarhou, 2007; von Economo & Koskinas, 1925) is ideally suited to identify biological gradients across the cortex because it is based on careful histological cytoarchitectural and myeloarchitectural cortical mapping. Using techniques that parcellate the cortex based upon microscopic changes in cell size, cell type, and myelin content at various sulcal depths allows for greater biologically interpretability of neuroimaging data across the cortex with differences captured across regions. Directly incorporating digitized atlases with *in-vivo* neuroimaging allows integration across biological scales of resolution.

The current study aims to combine the information provided from microscopy (utilizing the digitized vE-K atlas) with multi-modal MRI estimates of *in vivo* brain health (white matter diffusion and gray matter morphometry) to better refine our understanding of the biology underpinning *in vivo* neuroimaging and in turn, of aging brain structure. To elucidate the regional heterogeneity and heterochronicity in structural brain aging combining analysis of gray and white matter structural health within a histologically derived parcellation scheme, we model age-related trajectories of white matter diffusion paired with four-year longitudinal changes in gray matter morphometry to assess the degree to which these aging trajectories are chronologically linked, suggestive of overall neuronal degradation. We expect estimates of structural health to decline in tandem across the lifespan. Specifically, we expect heterogenous patterns in brain macrostructure to align with microstructurally defined parcellation schemes emphasizing the interconnected nature of neuronal subparts. Additionally, we predict that declines will be greatest in cortical regions associated with higher-order cognitive functioning, aligning with known patterns of neuronal layering and organization that underpin the sensorimotor-association gradient.

## 2 MATERIALS & METHODS

### 2.1 Participants

Participants were drawn from a healthy adult lifespan sample residing in the Dallas-Fort Worth Metroplex, the Dallas Area Longitudinal Lifespan Aging Study (DALLAS). All data were collected with participant’s informed consent and in accordance with institutional review board guidelines at The University of Texas at Dallas and the University of Texas Southwestern Medical Center. Participant recruitment was aimed towards collecting a representatively diverse sample of healthy adults with exclusions made for neurologic or psychiatric conditions, cardiovascular disease, head trauma with loss of consciousness > 5 minutes, diabetes, depression, dementia, substance abuse, and use of psychotropic medications. Depression was screened using the Center for Epidemiological Study Depression Scale (CES-D; Radloff, 1977) with a cutoff of ≤ 16, while dementia was screened using the Mini-Mental State Examination (MMSE; Folstein et al., 1975) with a cutoff of > 25. Additionally, all participants were required to be native English speakers (by the age of 6 yrs), right-handed, and with normal hearing and normal or corrected vision (at least 20/40). Finally, participants were excluded for any contraindications to MRI such as metallic implants or claustrophobia. These data were part of a larger, multi-wave longitudinal dataset collected between 2014 and 2020 that includes two separate neuroimaging sessions separated by an average of 50 months (approximately 4 years). In total, 105 participants successfully completed both waves of data collection including a full protocol of structural neuroimaging acquisitions. Of these 105 participants, three participants were removed due to brain abnormalities discovered in their neuroimaging data after completing the study. Therefore, the sample used was based upon the 102 participants that remained after completing quality assessment of the data (**Table 1** provides demographic information).

**Table 1.** Sample Demographics.

| Age group (range) | Sample size (M/F) | Age mean ± SD | Edu. mean ± SD | CESD mean ± SD | MMSE mean ± SD |
| --- | --- | --- | --- | --- | --- |
| YA (20-34) | 23 (8/15) | 27.00 ± 4.61 | 15.39 ± 2.21 | 4.91 ± 4.33 | 29.13 ± 1.01 |
| EM (35-54) | 23 (11/12) | 48.00 ± 4.78 | 15.04 ± 2.48 | 4.09 ± 4.25 | 29.09 ± 0.90 |
| LM (55-69) | 29 (11/18) | 61.10 ± 3.70 | 16.00 ± 2.39 | 4.41 ± 4.19 | 28.97 ± 0.78 |
| OA (70-86) | 27 (6/21) | 75.59 ± 5.03 | 16.01 ± 2.55 | 4.00 ± 3.91 | 28.89 ± 0.75 |
| **Total (20-86)** | **102 (36/66)** | **54.29 ± 18.25** | **15.66 ± 2.41** | **4.34 ± 4.12** | **29.01 ± 0.85** |

*Note.* Age was sampled and analyzed as a continuous variable in all analyses. Age groups are arbitrarily shown for sample visualization. YA – younger adults; EM – early middle-aged adults; LM – late middle-aged adults; OA – older adults. M – male; F – female; SD – standard deviation; Edu – education; CESD – Center for Epidemiological Studies – Depression scale; MMSE – Mini Mental Status Exam.

### 2.2 MRI Acquisition

Neuroimaging data were acquired on a single 3-Tesla Philips Achieva scanner with a 32-channel head coil using SENSE encoding (Philips Healthcare Systems, Best, Netherlands) at the University of Texas Southwestern’s Advanced Imaging Research Center. The scanning session included a T1-weighted acquisition and a single-shell diffusion-weighted imaging sequence to estimate various aspects of structural tissue health: T1-weighted (T1w) field echo MPRAGE images were collected with 160 sagittal slices, an isotropic voxel size of 1mm3, flip angle = 12°, TR/TE/TI = 8.1ms/3.7ms/1100ms, FOV = 204×256×160, matrix = 256×256, and a duration of 3:57min. Single-shell diffusion-weighted echo-planar imaging (EPI) sequence with 65 axial slices, a voxel size of 2×2×2.2 mm3 reconstructed to 0.85×0.85×2.2mm3, 30 diffusion directions at a b-value of 1000s/mm2 and 1 non-diffusion weighted at a b-value of 0 s/ mm2 (b0 image), TR/TE = 5608ms/51ms, FOV = 224×224×143, matrix = 112×112, and a duration of 4:19min.

### 2.3 Data Processing

T1w data were processed through an extensive suite of neuroimaging software packages to ensure quality and accuracy of the derived variables of interest. Raw data underwent manual quality assessment by trained researchers viewing multiple slices of each individual volume acquired for every participant to identify artifacts including anatomical abnormalities and participant movement distortions. Images were then processed through the automated *recon-all* function with iterative manual edits as part of the FreeSurfer package (Dale et al., 1999; Fischl & Dale, 2000). FreeSurfer allows calculation of surface-based estimates of volume, thickness, and surface-area measures as well as generation of pial and white matter surfaces. Each T1w image was processed in a cross-sectional fashion, in the absence of any information from the others waves of data from the same participant, and thus each participant has two separate and independent FreeSurfer reconstructions. A digitized version of the von Economo-Koskinas atlas was used to parcellate the brain based on its histological cytoarchitectural and laminar organization for each participant and each wave (Scholtens et al., 2018). Surface-based estimates of volume, thickness, and surface area were extracted from each parcellation of the vE-K atlas for each participant and each wave resulting in 43 regions in each hemisphere (**Figure 1**). See **Table 2** for list of regions and abbreviations.

**Figure 1.**
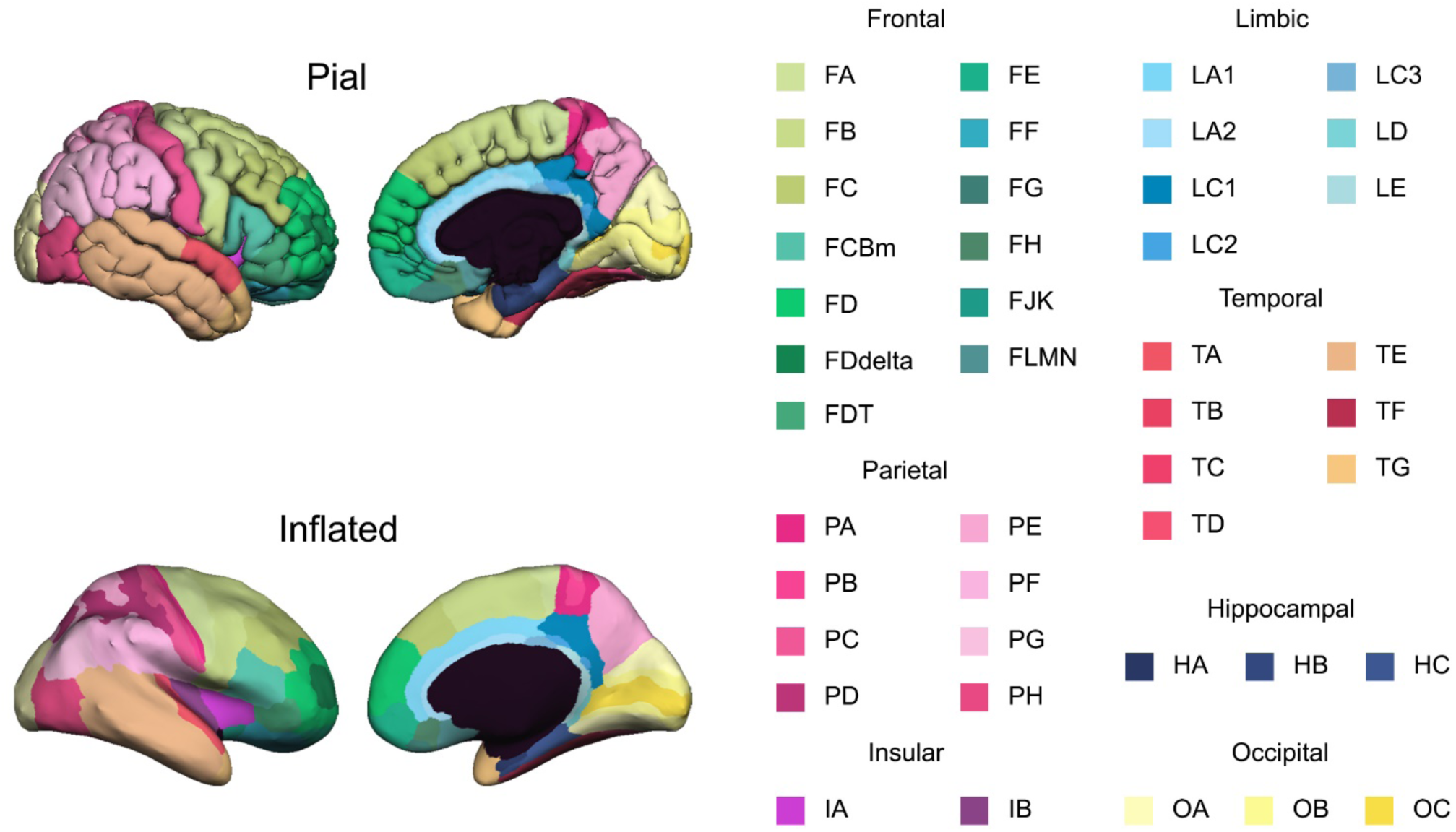
Von Economo-Koskinas atlas with the 43 digitized parcellated regions displayed on both the pial and inflated surfaces. Color scale shades correspond to lobar location: greens = frontal; dark blues = hippocampal; purples = insula; light blues = limbic; yellows = occipital; pinks = parietal; reds/tans = temporal cortices.

**Table 2.**
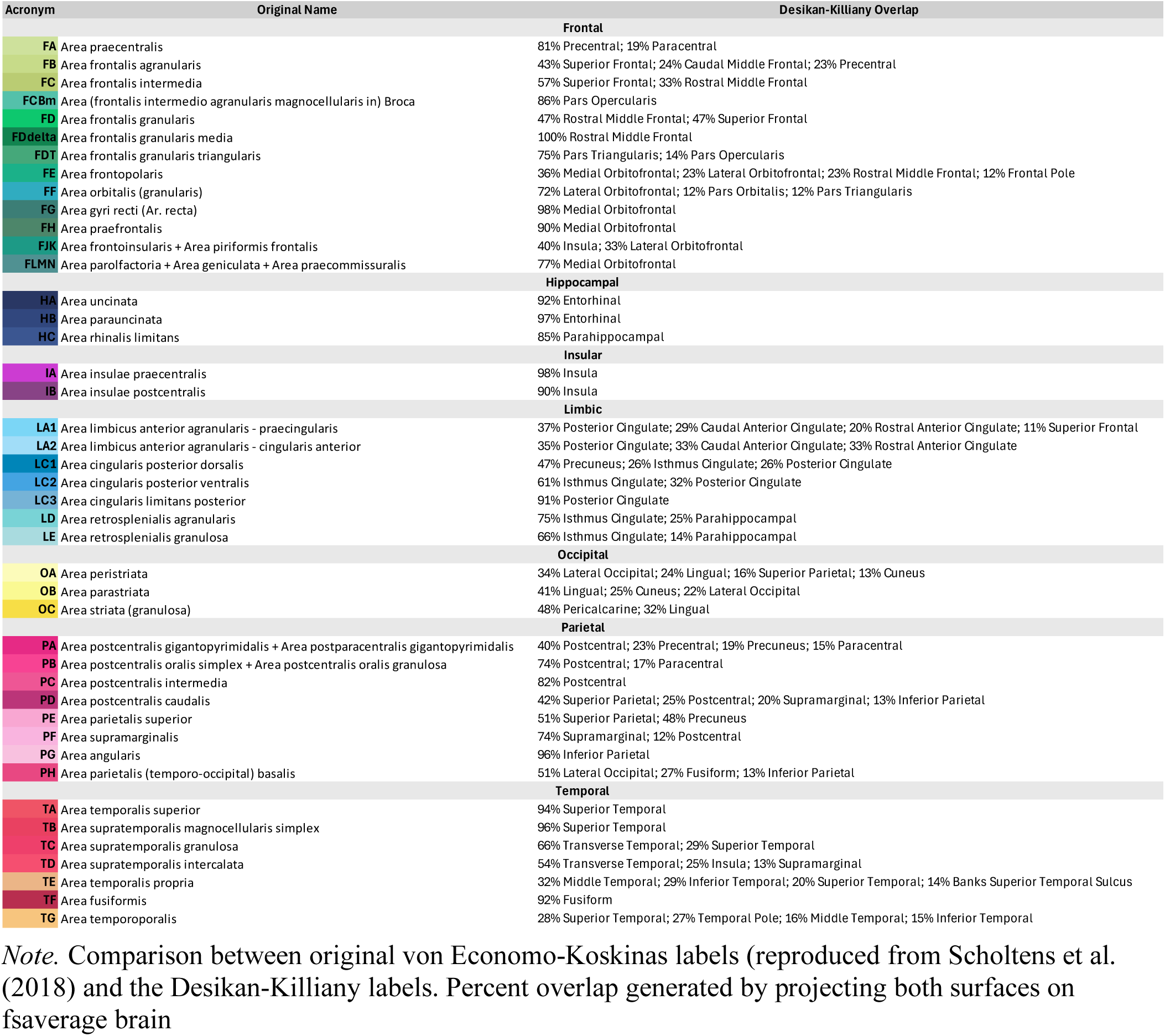
Von Economo-Koskinas region label, region name, and most closely corresponding regions in Desikan-Killiany label.

Raw diffusion weighted imaging data were quality assessed manually by trained researchers viewing multiple slices of each individual gradient acquired for every participant to identify artifacts including eddy current and intensity distortions, signal dropout, and participant movement distortions common to EPI imaging. Brain extraction was performed using FSL’s *bet* function on the non-diffusion weighted b0 image (Smith, 2002). Custom shell scripts and affine registration functions from the Advanced Normalization Tools (ANTs) software package (Avants et al., 2009) were used to apply the brain extraction mask to all gradients for increased accuracy in brain extraction across the entire acquisition. Automated preprocessing was performed using the software DTIprep (Liu et al., 2010) to account for any rotations applied to the gradients (Leemans & Jones, 2009). Diffusion tensor calculation was performed using the DSI Studio software package to obtain fractional anisotropy (FA), axial diffusivity (AD), radial diffusivity (RD), and mean diffusivity (MD) metrics (Yeh et al., 2013). To extract white matter metrics of interest from the wave 1 diffusion data, each parcellation from the vE-K atlas (**Figure 1**) was dilated by 3mm into the white matter of the wave 1 T1w image. A non-linear registration algorithm using the ANTs software program was used to align each participant’s wave 1 T1w MPRAGE scan to the wave 1 b0 image. This registration enabled mapping of the individual’s dilated parcellations of the vE-K atlas directly into diffusion space, resulting in extended vE-K parcel dilations into the white matter of the diffusion image (**Figure 2**). White matter metrics (FA, AD, RD, and MD) were extracted from this dilated space to ensure that the metrics were pulled from the voxels directly neighboring (within 3mm) the gray matter surface specific to each vE-K parcel. As our hypotheses of interest were not predicated on hemispheric differences, all variables were averaged within-modality across-hemispheres resulting in 43 gray matter and 43 adjacent white matter regions (**Figure 1**).

**Figure 2.**
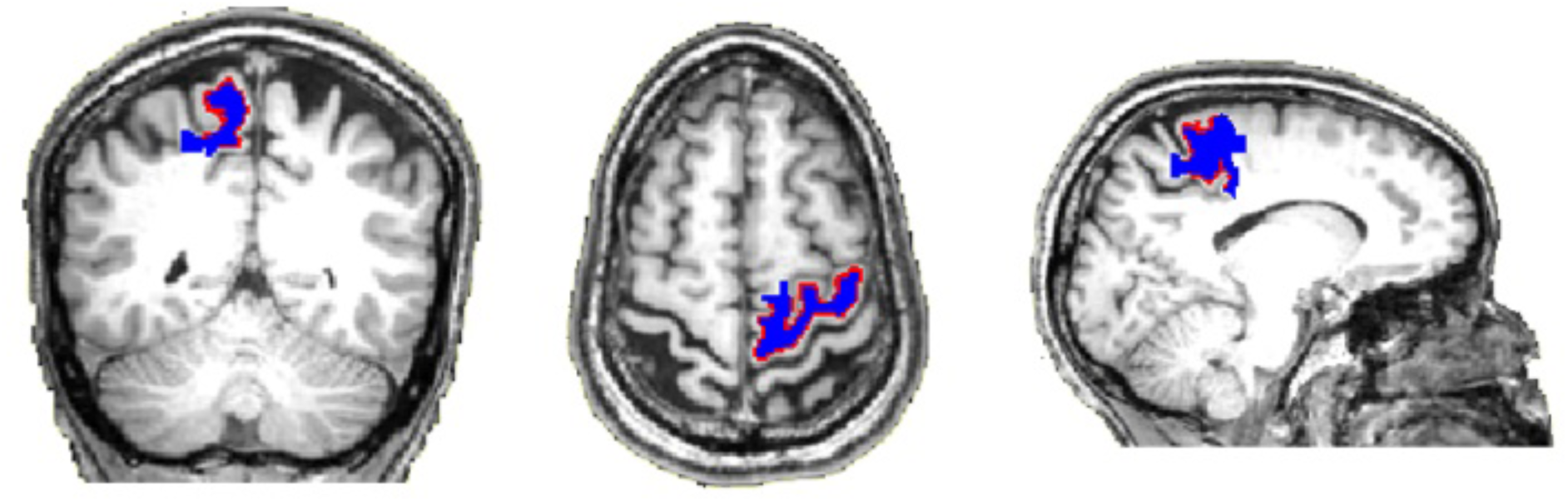
Coronal, axial, and sagittal views of an example participant’s brain displaying the “FA” region (precentral gyrus) from the Von Economo–Koskinas digital atlas. The border of the gray matter and white matter surface of the precentral region is outlined in red. White matter voxels included in a dilation of 3mm from the surface are in blue.

### 2.4 Statistical Analysis Approach

To estimate the associations of both within-subject and between-subjects proxies of structural health, linear mixed effects (LME) statistical models were employed to simultaneously account for all variables in a single model in a way that allows for modeling of higher-order interactions and accounts for both fixed and random effects. The full LME model can be broken down into two parts representing the fixed effects component (1) and the random effects component (2):

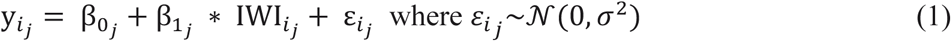

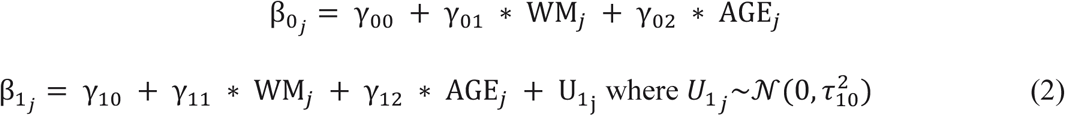

where *y* represents the gray matter morphometric feature across waves (i.e., volume, thickness or surface area); IWI is the inter-wave interval, or the time delay between a participant’s wave 1 and wave 2 MRI sessions; WM represents the white matter DTI metric extracted from the dilated regions at wave 1 (i.e., FA, RD, AD, MD); and AGE represents the participant’s age in years at their baseline MRI session at wave 1. Age, WM, and IWI were all mean-centered. Inherent to the hypotheses is examining how age and white matter diffusion differentially predict gray matter morphometric change. We used a single LME for each WM (FA, AD, MD, RD) and GM (thickness, volume, surface area) metric for each vE-K region and examined the main effects and interactions to understand the influence of each variable. Main effects included the change in gray matter predicted by the four-year inter-wave interval (IWI ∼ DGM), baseline age (Age ∼ DGM), and underlying WM (WM ∼ DGM). Additionally, the interaction of age and WM (Age x WM ∼ DGM) indicates how gray matter morphometry differences across the lifespan are differentially influenced by baseline white matter health proxies. Importantly, because all variables and interactions are included in the LME model, focusing on specific main effects or interactions accounts for other variables (similar to considering them as regressors or covariates). All statistical tests were run using the “lmerTest” Linear Mixed-Effects Models using ‘Eigen’ and S4 (Bates et al., 2015) library within the R statistics software package (R Core Team, 2021).

## 3 RESULTS

### 3.1 Analytic Overview

Linear mixed effects models returned main effects and interactions for each of the terms of IWI, baseline white matter diffusion, and baseline age on the change in gray matter morphometry across the regions. Results from age and white matter terms (and their interactions) on change in gray matter morphometry were informative and will be discussed below, however the interactions with IWI were non-significant across the majority of cortical regions, which would be expected with a tight follow-up range. For this reason, we report these interactions only in *Supplemental Material (Tables S5-7, Figures S5-7)*. The four sets of LME results across the models are summarized in **Tables 3-6** and are plotted on the inflated cortical surface in corresponding panels A-D of **Figure 3**. Tables are organized into two halves with the fractional anisotropy results on the left and mean diffusivity results on the right (results for radial and axial diffusivities are presented in *Supplemental Tables 1-4* and *Supplemental Figures 1-4*). Columns represent the three gray matter morphometric features (cortical thickness, volume, and surface area), while rows are the vE-K regions sorted by lobe. Color coding matches the region legend in **Figure 1**.

**Figure 3:**
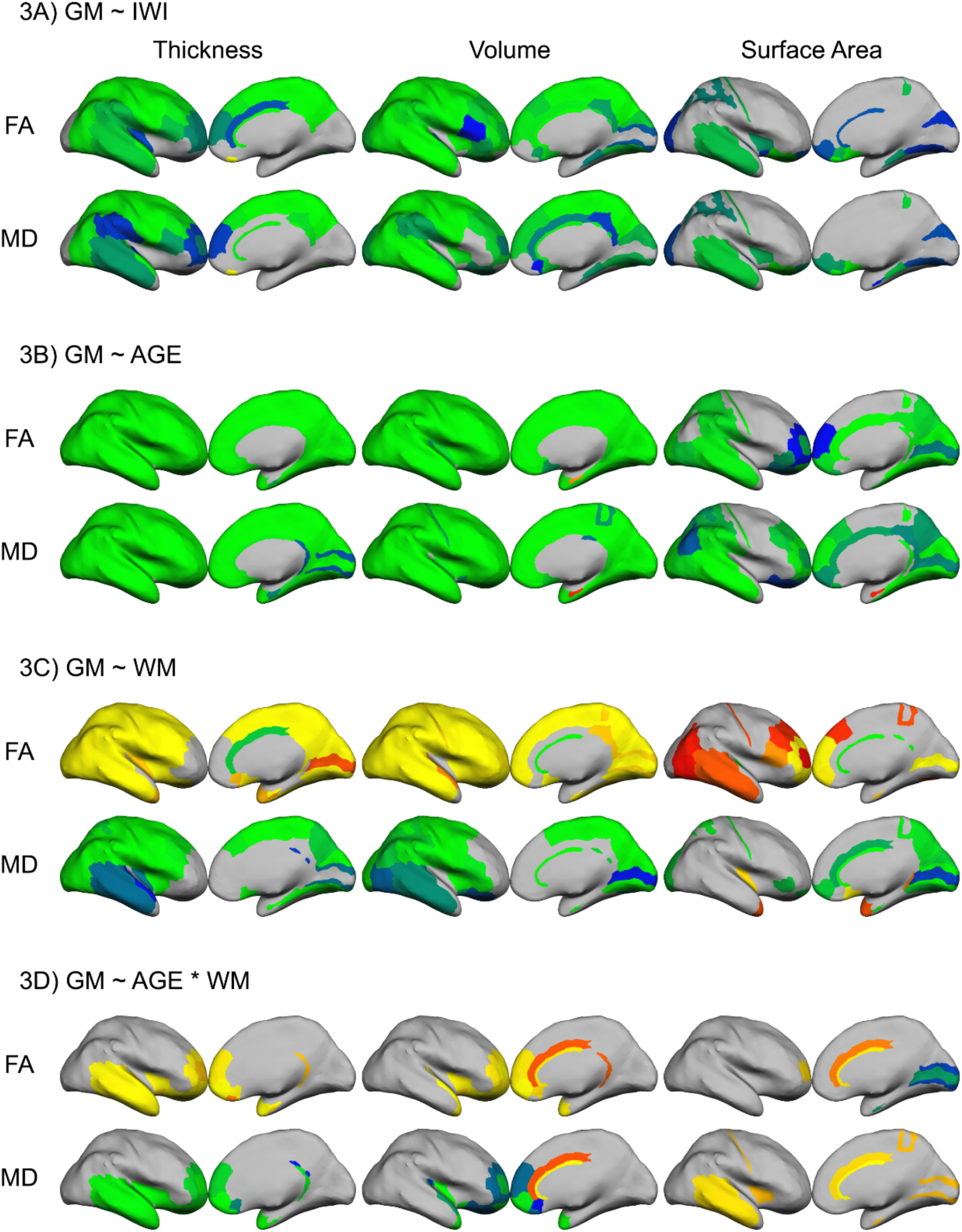
Regional main effects of A) inter-wave-interval, B) age, C) white matter metrics, on gray matter morphometry and D) interaction effects of age and white matter metrics, plotted on the inflated surface using von Economo – Koskinas atlas.

### 3.2 Change in gray matter morphometry across the inter-wave interval

Across the roughly 4-year IWI, we found significant gray matter change in all three measures of cortical morphometry (**Figure 3A**; **Table 3**). Amongst these, longitudinal decreases in volume were the most robust and were observed across the majority of regions except for primary occipital (OC), hippocampal (HA and HB), posterior limbic (LC3, LD, and LE), temporopolar (TG), and various frontopolar (FE) and orbitofrontal regions (FJK and FG). The most significant declines in volume were seen in prerolandic frontal, parietal, and some temporal regions. Changes in thickness were less robust, but still found to be significant in about half of the cortical regions. Areas closest to the pre- and post-central gyri underwent the most significant change in thickness over time, while occipital and orbitofrontal areas were non-significant. Surface area was the most resilient to aging with only a handful of areas showing significant longitudinal decreases in surface area, with many non-significant regions mirroring the thickness results, such as the orbitofrontal areas and some occipital regions. Additionally, middle temporal (TE), auditory areas (TB), and parts of the post-central (PB and PD) were among the few areas demonstrating significant 4-year decline across all three aspects of morphometry. These opposing regional thickness-surface area findings suggest there are fundamental differences in these two aspects of morphometry. Parts of the frontopolar and orbitofrontal regions evidence significant decreases in surface area and volume but increases in thickness (FG, FF, FE, and FJK). While this could be an example of differential aging of each morphometric feature in these regions, it is also likely that there are inconsistencies in estimates of thickness from these posterior frontal regions that are susceptible to imaging artifacts.

**Table 3.** Gray matter change associated with 4-year inter-wave-interval.

| Gray matter change associated with 4-year inter-wave-interval |  |  |  |  |  |  |  |  |  |  |  |  |
| --- | --- | --- | --- | --- | --- | --- | --- | --- | --- | --- | --- | --- |
| VE | FA |  |  |  |  |  | MD |  |  |  |  |  |
|  | Thickness |  | Volume |  | Surface Area |  | Thickness |  | Volume |  | Surface Area |  |
|  | t-value | p-value | t-value | p-value | t-value | p-value | t-value | p-value | t-value | p-value | t-value | p-value |
| FA | -4.5655 | 8.8E-06 | -4.7877 | 3.3E-06 | -1.6697 | 0.0966 | -4.2896 | 2.8E-05 | -4.3366 | 2.3E-05 | -1.6121 | 0.1085 |
| FB | -3.8217 | 0.0002 | -3.3229 | 0.0011 | -1.0774 | 0.2826 | -3.9336 | 0.0001 | -3.4547 | 0.0007 | -1.2817 | 0.2015 |
| FC | -2.9291 | 0.0038 | -2.6511 | 0.0087 | -1.0915 | 0.2764 | -2.9472 | 0.0036 | -2.7173 | 0.0072 | -1.2532 | 0.2116 |
| FCBm | -2.5241 | 0.0124 | -1.9907 | 0.0479 | -0.8562 | 0.3929 | -2.3744 | 0.0185 | -1.7928 | 0.0746 | -0.7707 | 0.4418 |
| FD | -2.3017 | 0.0224 | -2.9807 | 0.0032 | -1.6194 | 0.1070 | -2.0834 | 0.0385 | -2.6898 | 0.0078 | -1.5712 | 0.1177 |
| FDdelta | -2.2761 | 0.0239 | -2.5529 | 0.0114 | -1.0261 | 0.3061 | -2.2924 | 0.0229 | -2.2763 | 0.0239 | -0.8791 | 0.3804 |
| FDT | -1.7625 | 0.0795 | -2.4535 | 0.0150 | -1.4725 | 0.1425 | -1.3085 | 0.1922 | -1.8421 | 0.0670 | -1.1756 | 0.2412 |
| FE | 1.1887 | 0.2360 | -0.3354 | 0.7377 | -2.1345 | 0.0340 | 1.7436 | 0.0828 | 0.0096 | 0.9923 | -2.3532 | 0.0196 |
| FF | 1.1758 | 0.2411 | -2.5944 | 0.0102 | -4.0514 | 7.3E-05 | 1.4569 | 0.1467 | -2.3674 | 0.0189 | -4.1279 | 5.4E-05 |
| FG | 4.2101 | 3.9E-05 | -0.1955 | 0.8452 | -3.5556 | 0.0005 | 4.0050 | 8.8E-05 | -0.0712 | 0.9433 | -3.3266 | 0.0010 |
| FH | 0.1279 | 0.8983 | -2.4052 | 0.0171 | -2.4716 | 0.0143 | 0.5047 | 0.6143 | -1.9971 | 0.0472 | -2.3731 | 0.0186 |
| FJK | 0.2333 | 0.8158 | -0.5064 | 0.6131 | -2.0636 | 0.0404 | 0.7447 | 0.4574 | 0.0925 | 0.9264 | -1.5794 | 0.1158 |
| FLMN | -0.0980 | 0.9220 | -2.8376 | 0.0050 | -3.1960 | 0.0016 | 0.3687 | 0.7127 | -2.6478 | 0.0088 | -3.5736 | 0.0004 |
| HA | -0.2785 | 0.7810 | -1.2135 | 0.2264 | -1.3493 | 0.1788 | -0.2828 | 0.7776 | -1.2247 | 0.2221 | -1.2030 | 0.2304 |
| HB | 0.6999 | 0.4848 | -1.6170 | 0.1075 | -1.8970 | 0.0593 | 0.6728 | 0.5019 | -1.7444 | 0.0827 | -1.9941 | 0.0475 |
| HC | -1.2589 | 0.2096 | -2.2792 | 0.0237 | -2.3381 | 0.0204 | -1.2491 | 0.2131 | -2.3416 | 0.0202 | -2.5125 | 0.0128 |
| IA | -1.4284 | 0.1548 | -3.2457 | 0.0014 | -2.3558 | 0.0195 | -1.0662 | 0.2877 | -2.8959 | 0.0042 | -2.4345 | 0.0158 |
| IB | -2.1053 | 0.0365 | -1.6071 | 0.1096 | -1.1922 | 0.2346 | -1.8698 | 0.0630 | -1.3534 | 0.1775 | -1.0957 | 0.2746 |
| LA1 | -2.1230 | 0.0350 | -2.9279 | 0.0038 | -1.8721 | 0.0627 | -1.5584 | 0.1208 | -2.2619 | 0.0248 | -1.6636 | 0.0978 |
| LA2 | -4.4905 | 1.2E-05 | -4.3546 | 2.1E-05 | -2.1387 | 0.0337 | -3.1290 | 0.0020 | -2.9573 | 0.0035 | -1.6609 | 0.0983 |
| LC1 | -3.8061 | 0.0002 | -2.3763 | 0.0185 | -0.3857 | 0.7002 | -2.7780 | 0.0060 | -2.0807 | 0.0388 | -0.6681 | 0.5048 |
| LC2 | -3.8664 | 0.0002 | -2.7589 | 0.0063 | -1.0466 | 0.2966 | -3.4651 | 0.0007 | -2.9142 | 0.0040 | -1.2449 | 0.2147 |
| LC3 | -1.1004 | 0.2725 | -1.8020 | 0.0731 | -1.2368 | 0.2176 | 0.4586 | 0.6471 | -1.0457 | 0.2970 | -1.2872 | 0.1996 |
| LD | -1.6892 | 0.0928 | -1.9176 | 0.0566 | -0.4748 | 0.6355 | -1.7167 | 0.0876 | -1.9192 | 0.0564 | -0.4588 | 0.6469 |
| LE | -0.7087 | 0.4794 | -0.9969 | 0.3200 | -1.1774 | 0.2405 | -0.4621 | 0.6445 | -0.6400 | 0.5229 | -1.1074 | 0.2695 |
| OA | -1.0974 | 0.2738 | -2.6337 | 0.0091 | -2.0537 | 0.0413 | -0.8573 | 0.3923 | -2.5050 | 0.0131 | -2.1213 | 0.0352 |
| OB | 0.0948 | 0.9246 | -2.1588 | 0.0321 | -1.5463 | 0.1237 | -0.2429 | 0.8083 | -2.4763 | 0.0141 | -1.6285 | 0.1050 |
| OC | -1.2528 | 0.2118 | -1.6852 | 0.0935 | -1.1109 | 0.2680 | -0.7885 | 0.4313 | -1.4431 | 0.1506 | -1.1496 | 0.2517 |
| PA | -4.6937 | 5.0E-06 | -3.9983 | 9.0E-05 | -1.5661 | 0.1189 | -3.9772 | 9.8E-05 | -3.3649 | 0.0009 | -1.5252 | 0.1288 |
| PB | -4.0425 | 7.6E-05 | -4.5186 | 1.1E-05 | -2.5112 | 0.0128 | -3.6755 | 0.0003 | -4.4262 | 1.6E-05 | -2.6224 | 0.0094 |
| PC | -3.3243 | 0.0011 | -3.4170 | 0.0008 | -1.0287 | 0.3049 | -3.0563 | 0.0026 | -3.1957 | 0.0016 | -1.0517 | 0.2942 |
| PD | -4.8406 | 2.6E-06 | -4.1389 | 5.2E-05 | -2.2816 | 0.0236 | -4.4615 | 1.4E-05 | -3.9329 | 0.0001 | -2.2888 | 0.0232 |
| PE | -4.3513 | 2.2E-05 | -4.0340 | 7.9E-05 | -1.4146 | 0.1588 | -4.0427 | 7.6E-05 | -3.9610 | 0.0001 | -1.5415 | 0.1248 |
| PF | -2.5184 | 0.0126 | -2.8765 | 0.0045 | -1.9002 | 0.0589 | -2.0693 | 0.0398 | -2.4144 | 0.0167 | -1.7226 | 0.0865 |
| PG | -3.0985 | 0.0022 | -3.4272 | 0.0007 | -1.6789 | 0.0948 | -3.0852 | 0.0023 | -3.2984 | 0.0012 | -1.7512 | 0.0815 |
| PH | -3.1599 | 0.0018 | -3.1457 | 0.0019 | -1.3834 | 0.1681 | -3.1991 | 0.0016 | -3.0819 | 0.0024 | -1.3994 | 0.1633 |
| TA | -3.3845 | 0.0009 | -2.9197 | 0.0039 | -1.2621 | 0.2084 | -3.2916 | 0.0012 | -2.6784 | 0.0080 | -1.1160 | 0.2658 |
| TB | -3.2421 | 0.0014 | -3.0600 | 0.0025 | -2.1864 | 0.0300 | -3.3057 | 0.0011 | -3.0010 | 0.0030 | -2.1700 | 0.0312 |
| TC | -2.0646 | 0.0403 | -2.1686 | 0.0313 | -1.1426 | 0.2546 | -1.9980 | 0.0471 | -1.8967 | 0.0593 | -0.8323 | 0.4063 |
| TD | -2.2116 | 0.0282 | -1.2701 | 0.2055 | -0.2444 | 0.8072 | -1.6538 | 0.0998 | -1.2652 | 0.2073 | -0.4904 | 0.6244 |
| TE | -2.6240 | 0.0094 | -3.9960 | 9.1E-05 | -2.5073 | 0.0130 | -2.3441 | 0.0201 | -3.7156 | 0.0003 | -2.6014 | 0.0100 |
| TF | -1.1779 | 0.2403 | -1.6953 | 0.0916 | -1.3191 | 0.1887 | -1.1339 | 0.2582 | -1.7670 | 0.0788 | -1.5865 | 0.1142 |
| TG | -0.0294 | 0.9766 | -0.1127 | 0.9104 | -1.3236 | 0.1872 | 0.0944 | 0.9249 | -0.0916 | 0.9271 | -1.4244 | 0.1559 |

Note. Regional linear mixed effects model results from figure 3A. Green to red cell shading indicates magnitude, green text color indicates significance at $p < .05$ .

### 3.3 Change in gray matter morphometry associated with age

Longitudinal changes in gray matter morphometry were found to be highly associated with participant’s age such that increasing age is associated with more gray matter loss over 4 years. There was a significant main effect of declining gray matter thickness and volume across the lifespan in almost every region analyzed (**Table 4**, **Figure 3B**). Frontal, posterior parietal, and temporal regions demonstrated the most significant change in thickness and volume across the lifespan. Age-related surface area declines were also prevalent in almost half of all regions analyzed, with effects primarily in the pre-frontal, posterior parietal, lateral temporal, anterior limbic, and occipital regions (**Table 4**, **Figure 3B**). The ubiquity of these significant findings warrants a closer examination of the gradients of effects across the cortex to better understand the emerging patterns characterizing lifespan change and are discussed in section 4.2.

**Table 4.** Gray matter change associated with age.

Gray matter change associated with age
| VE | FA |  |  |  |  |  |  | MD |  |  |  |  |  |
| --- | --- | --- | --- | --- | --- | --- | --- | --- | --- | --- | --- | --- | --- |
|  | Thickness |  | Volume |  | Surface Area |  |  | Thickness |  | Volume |  | Surface Area |  |
|  | t-value | p-value | t-value | p-value | t-value | p-value |  | t-value | p-value | t-value | p-value | t-value | p-value |
| FA | -9.6115 | 3.6E-18 | -7.9560 | 1.4E-13 | -1.0564 | 0.2921 |  | -6.6349 | 3.1E-10 | -5.4937 | 1.2E-07 | -1.0735 | 0.2844 |
| FB | -7.7533 | 4.8E-13 | -5.6503 | 5.6E-08 | -1.7821 | 0.0763 |  | -8.7145 | 1.2E-15 | -6.0582 | 6.9E-09 | -1.7082 | 0.0892 |
| FC | -7.8472 | 2.7E-13 | -4.4026 | 1.8E-05 | -1.2448 | 0.2147 |  | -8.9565 | 2.6E-16 | -6.3746 | 1.3E-09 | -2.5909 | 0.0103 |
| FCBm | -8.3257 | 1.4E-14 | -3.5635 | 0.0005 | -0.0617 | 0.9508 |  | -9.5973 | 4.0E-18 | -4.9403 | 1.7E-06 | -0.6199 | 0.5360 |
| FD | -9.9519 | 3.8E-19 | -7.3234 | 6.1E-12 | -1.9859 | 0.0484 |  | -8.8723 | 4.5E-16 | -8.0450 | 8.1E-14 | -3.1049 | 0.0022 |
| FDdelta | -7.3869 | 4.2E-12 | -6.1027 | 5.5E-09 | -2.4702 | 0.0144 |  | -6.9966 | 4.1E-11 | -5.7542 | 3.3E-08 | -2.4320 | 0.0159 |
| FDT | -10.6772 | 2.9E-21 | -8.3730 | 1.1E-14 | -2.3636 | 0.0191 |  | -9.9109 | 5.0E-19 | -7.2217 | 1.1E-11 | -1.4126 | 0.1594 |
| FE | -6.6769 | 2.5E-10 | -7.2674 | 8.5E-12 | -2.7982 | 0.0057 |  | -6.6431 | 3.0E-10 | -6.8854 | 7.6E-11 | -2.3291 | 0.0209 |
| FF | -7.5354 | 1.8E-12 | -7.3852 | 4.3E-12 | -2.2321 | 0.0267 |  | -6.6224 | 3.3E-10 | -6.4057 | 1.1E-09 | -2.0784 | 0.0390 |
| FG | -3.2979 | 0.0012 | -6.3520 | 1.5E-09 | -2.5064 | 0.0130 |  | -3.4202 | 0.0008 | -6.2919 | 2.0E-09 | -2.4876 | 0.0137 |
| FH | -4.1316 | 5.3E-05 | -5.0505 | 1.0E-06 | -1.0761 | 0.2832 |  | -4.0431 | 7.6E-05 | -4.2985 | 2.7E-05 | -0.7590 | 0.4488 |
| FJK | -6.7690 | 1.5E-10 | -3.8502 | 0.0002 | 0.0705 | 0.9439 |  | -4.2529 | 3.3E-05 | -2.3796 | 0.0183 | -0.4925 | 0.6229 |
| FLMN | -3.9209 | 0.0001 | -2.3995 | 0.0174 | -0.0013 | 0.9990 |  | -3.3972 | 0.0008 | -2.7744 | 0.0061 | -1.3014 | 0.1946 |
| HA | -1.4104 | 0.1600 | 2.3265 | 0.0210 | 1.7108 | 0.0887 |  | -2.2864 | 0.0233 | 2.0403 | 0.0427 | 2.0319 | 0.0435 |
| HB | -2.4458 | 0.0153 | -2.7074 | 0.0074 | -1.3197 | 0.1885 |  | -2.2738 | 0.0241 | -3.2458 | 0.0014 | -1.8280 | 0.0691 |
| HC | -3.6910 | 0.0003 | -5.4718 | 1.4E-07 | -4.2361 | 3.5E-05 |  | -2.8352 | 0.0051 | -5.7820 | 2.9E-08 | -5.6241 | 6.4E-08 |
| IA | -6.6059 | 3.6E-10 | -7.3891 | 4.2E-12 | -0.9628 | 0.3368 |  | -4.2782 | 2.9E-05 | -5.1132 | 7.5E-07 | -1.5875 | 0.1140 |
| IB | -5.3341 | 2.6E-07 | -2.7204 | 0.0071 | 0.8366 | 0.4039 |  | -3.8966 | 0.0001 | -2.9289 | 0.0038 | -0.5648 | 0.5729 |
| LA1 | -8.6041 | 2.5E-15 | -7.4864 | 2.3E-12 | -3.8468 | 0.0002 |  | -7.5533 | 1.6E-12 | -5.4971 | 1.2E-07 | -2.3870 | 0.0179 |
| LA2 | -3.0762 | 0.0024 | -5.5416 | 9.6E-08 | -3.9868 | 9.4E-05 |  | -3.0283 | 0.0028 | -1.5817 | 0.1153 | 0.1541 | 0.8777 |
| LC1 | -7.6210 | 1.1E-12 | -6.0919 | 5.8E-09 | -2.8634 | 0.0046 |  | -7.0180 | 3.6E-11 | -5.6547 | 5.5E-08 | -2.2838 | 0.0235 |
| LC2 | -9.6830 | 2.3E-18 | -4.8243 | 2.8E-06 | -1.6152 | 0.1079 |  | -7.7023 | 6.5E-13 | -4.0972 | 6.1E-05 | -1.5364 | 0.1261 |
| LC3 | -4.1306 | 5.4E-05 | -3.5341 | 0.0005 | -0.9115 | 0.3632 |  | -3.2361 | 0.0014 | -2.1991 | 0.0290 | -0.2185 | 0.8273 |
| LD | -5.3331 | 2.6E-07 | -2.7625 | 0.0063 | -0.3026 | 0.7625 |  | -5.1107 | 7.6E-07 | -2.8099 | 0.0055 | -0.4882 | 0.6259 |
| LE | -3.1301 | 0.0020 | -3.8428 | 0.0002 | -3.1648 | 0.0018 |  | -2.1745 | 0.0309 | -3.4383 | 0.0007 | -3.0807 | 0.0024 |
| OA | -7.4989 | 2.2E-12 | -5.3974 | 1.9E-07 | -3.3303 | 0.0010 |  | -7.5005 | 2.2E-12 | -5.3244 | 2.8E-07 | -3.3001 | 0.0011 |
| OB | -3.1989 | 0.0016 | -4.3113 | 2.6E-05 | -3.6757 | 0.0003 |  | -2.1373 | 0.0338 | -4.6288 | 6.7E-06 | -4.6346 | 6.5E-06 |
| OC | -5.0500 | 1.0E-06 | -3.7623 | 0.0002 | -2.2504 | 0.0255 |  | -4.9667 | 1.5E-06 | -3.7436 | 0.0002 | -2.2858 | 0.0233 |
| PA | -6.4968 | 6.6E-10 | -3.3636 | 0.0009 | 0.8371 | 0.4035 |  | -5.7808 | 2.9E-08 | -2.3406 | 0.0203 | 1.4707 | 0.1430 |
| PB | -4.0577 | 7.2E-05 | -5.5916 | 7.5E-08 | -3.8809 | 0.0001 |  | -3.6016 | 0.0004 | -4.9741 | 1.4E-06 | -3.4475 | 0.0007 |
| PC | -8.7666 | 8.8E-16 | -7.5852 | 1.3E-12 | -1.3384 | 0.1823 |  | -7.1884 | 1.3E-11 | -6.6157 | 3.4E-10 | -1.6613 | 0.0983 |
| PD | -7.4737 | 2.5E-12 | -4.9117 | 1.9E-06 | -2.7601 | 0.0063 |  | -6.8686 | 8.4E-11 | -4.4587 | 1.4E-05 | -2.5344 | 0.0120 |
| PE | -9.3518 | 2.0E-17 | -7.6715 | 7.8E-13 | -2.6908 | 0.0077 |  | -9.1405 | 8.0E-17 | -7.3952 | 4.0E-12 | -2.3447 | 0.0200 |
| PF | -9.1102 | 9.7E-17 | -6.6197 | 3.4E-10 | -2.8070 | 0.0055 |  | -7.9693 | 1.3E-13 | -6.1532 | 4.2E-09 | -2.9590 | 0.0035 |
| PG | -9.4858 | 8.4E-18 | -6.9216 | 6.2E-11 | -1.5698 | 0.1181 |  | -9.2207 | 4.7E-17 | -7.0764 | 2.6E-11 | -2.1564 | 0.0323 |
| PH | -10.2781 | 4.3E-20 | -6.5058 | 6.3E-10 | -2.6324 | 0.0092 |  | -9.7291 | 1.7E-18 | -6.2329 | 2.8E-09 | -2.8784 | 0.0044 |
| TA | -12.1458 | 1.1E-25 | -7.0480 | 3.0E-11 | 0.1397 | 0.8890 |  | -10.9657 | 4.0E-22 | -6.3447 | 1.5E-09 | -0.3957 | 0.6928 |
| TB | -10.2051 | 7.0E-20 | -6.2727 | 2.2E-09 | -0.0622 | 0.9505 |  | -9.5489 | 5.5E-18 | -6.2293 | 2.8E-09 | -0.6692 | 0.5042 |
| TC | -8.1851 | 3.4E-14 | -4.2658 | 3.1E-05 | 0.1456 | 0.8844 |  | -7.9407 | 1.5E-13 | -3.7613 | 0.0002 | 0.5873 | 0.5577 |
| TD | -7.6048 | 1.2E-12 | -2.3622 | 0.0191 | 0.4642 | 0.6430 |  | -6.1846 | 3.6E-09 | -1.8746 | 0.0623 | 0.2788 | 0.7807 |
| TE | -10.3960 | 1.9E-20 | -8.3194 | 1.5E-14 | -3.0500 | 0.0026 |  | -8.8757 | 4.4E-16 | -7.7614 | 4.5E-13 | -3.8934 | 0.0001 |
| TF | -5.5792 | 8.0E-08 | -5.1241 | 7.1E-07 | -2.6836 | 0.0079 |  | -5.6864 | 4.7E-08 | -5.9467 | 1.2E-08 | -3.3587 | 0.0009 |
| TG | -6.8442 | 9.6E-11 | -7.4455 | 3.0E-12 | -1.3150 | 0.1901 |  | -6.1353 | 4.6E-09 | -6.9833 | 4.4E-11 | -1.8950 | 0.0596 |

Note. Regional linear mixed effects model results from figure 3B. Green to red cell shading indicates magnitude, green text color indicates significance at $p < .05$ .

An overall gradient of association strength is evident across regions throughout the cortex. Changes in the relationship between thickness or volume with age attenuates when moving from higher order association-like cortices to primary motor and sensory areas. This is most evident in hippocampal, occipital, and insular areas that all have weak associations. Primary sensory areas demonstrate weak associations within the parietal lobe, strengthening when moving posteriorly towards angular (PG) and supramarginal gyri (PF), and in the temporal lobe, strengthening when moving away from transverse temporal (TC and TD) and fusiform areas (TF) toward inferior and middle temporal (TE) and temporopolar (TG). The frontal lobe shows a less clear gradient. While triangular gyrus (FDT), middle frontal gyrus (FDT), orbital frontal (FF) and frontal pole (FE) areas demonstrate consistently high age-related change, some posterior motor areas also show strong morphometric differences. Surface area measures show differential age-related relationships that are much weaker overall than both thickness and volume. Interestingly, most of the association cortices exhibiting the greatest change in thickness and volume, were also significant with surface area, albeit to a lesser degree. This includes much of the prefrontal, inferior parietal, and inferior temporal regions, while prerolandic frontal regions surrounding motor areas and supratemporal regions surrounding auditory areas remained non-significant.

Very few areas showed non-significant changes (areas in gray) or increases (areas in warm color scale) in morphometry with age. Hippocampal regions showed some of the weakest associations, even showing age-related increases in volume and surface area in the entorhinal (HA). Similarly, primary areas of the parietal and temporal lobes, specifically the precentral (PA) and supratemporal regions (TA, TC, and TD) show increases in surface area, albeit non-significant.

We also note that because diffusivity estimates were included in the omnibus models, when isolating the main effects of age in the LMEs, these proxies of white matter health were essentially treated as regressors. Comparing across models there are differences between fractional anisotropy and mean diffusivity that suggest a differential influence in aging by accounting for each (**Table 4**). In general, declines in thickness and volume were influenced more when accounting for FA than MD. Regionally, accounting for FA led to more significant age-related change in the precentral (FA) compared to accounting for MD, whereas accounting for MD leads to more significant age-related change in the agranular and intermediate frontal areas (FB and FC) than when accounting for FA. Differences in the change in surface area were primarily in limbic regions, as accounting for FA led to much stronger declines in surface area than accounting for MD, especially in anterior limbic (LA2). Other important differences were observed between aspects of thickness and volume. Overall, greater age-related changes were observed when modeling thickness. Thickness was disproportionately influencing estimates from supratemporal regions leading to much stronger relationships with age than volume. Interestingly, areas that showed more declines in thickness compared to volume were also the areas that showed no significant differences in surface area with age.

### 3.4 Change in gray matter morphometry associated with baseline measures of white matter diffusion

One major aim of this study was to identify areas in which white matter structure was significantly coupled with longitudinal changes in gray matter morphometry. Overall, large portions of the cortex show a significant gray matter – white matter coupling, suggesting that there are biological characteristics linking the two tissue types. Specifically, baseline white matter FA and MD were found to significantly influence the change in gray matter morphometry such that poorer white matter health (decreased FA or increased diffusivity) was associated with greater morphometric declines. Regionally, associations were strongest in the sensory and motor areas of the frontal, parietal, and temporal cortices, primarily in the areas encompassing the pre and postcentral gyri (**Table 5**, **Figure 3C**). While results were consistent across regional models, there were several gradients of effects and differences across measures that were notable, and arediscussed in detail in section 4.3.

**Table 5.** Gray matter change associated with baseline white matter.

Gray matter change associated with baseline white matter
| VE | FA |  |  |  |  |  | MD |  |  |  |  |  |
| --- | --- | --- | --- | --- | --- | --- | --- | --- | --- | --- | --- | --- |
|  | Thickness |  | Volume |  | Surface Area |  | Thickness |  | Volume |  | Surface Area |  |
|  | t-value | p-value | t-value | p-value | t-value | p-value | t-value | p-value | t-value | p-value | t-value | p-value |
| FA | 8.7061 | 1.3E-15 | 6.3018 | 1.9E-09 | -0.7950 | 0.4276 | -8.3441 | 1.3E-14 | -5.5542 | 9.0E-08 | 0.5452 | 0.5862 |
| FB | 6.0971 | 5.6E-09 | 4.4794 | 1.3E-05 | 0.9985 | 0.3193 | -4.6780 | 5.4E-06 | -3.2842 | 0.0012 | -0.8330 | 0.4059 |
| FC | 4.0081 | 8.7E-05 | 4.2239 | 3.7E-05 | 2.0499 | 0.0417 | -2.9631 | 0.0034 | -1.0200 | 0.3090 | 0.0329 | 0.9738 |
| FCBm | 5.1124 | 7.5E-07 | 5.1850 | 5.4E-07 | 2.3234 | 0.0212 | -3.3094 | 0.0011 | -2.8771 | 0.0045 | -1.5139 | 0.1317 |
| FD | 0.7592 | 0.4487 | 4.7868 | 3.3E-06 | 3.6170 | 0.0004 | -1.1068 | 0.2697 | -0.6156 | 0.5389 | -0.3746 | 0.7084 |
| FDdelta | 1.2819 | 0.2014 | 2.9228 | 0.0039 | 1.9786 | 0.0493 | -0.0416 | 0.9669 | -0.0841 | 0.9331 | -0.2339 | 0.8153 |
| FDT | 1.8774 | 0.0619 | 3.1461 | 0.0019 | 1.7413 | 0.0832 | -1.1426 | 0.2546 | -3.2447 | 0.0014 | -2.5460 | 0.0117 |
| FE | 0.6009 | 0.5486 | 3.7069 | 0.0003 | 3.0475 | 0.0026 | 0.9431 | 0.3468 | -1.5785 | 0.1161 | -2.6256 | 0.0093 |
| FF | 3.8165 | 0.0002 | 4.2035 | 4.0E-05 | 0.6901 | 0.4910 | -1.9666 | 0.0506 | -2.1504 | 0.0327 | -0.6234 | 0.5337 |
| FG | 1.9377 | 0.0541 | -0.2421 | 0.8090 | -1.7691 | 0.0784 | -0.2519 | 0.8014 | 1.2403 | 0.2163 | 1.5332 | 0.1268 |
| FH | 2.4949 | 0.0134 | 1.3489 | 0.1789 | -0.6908 | 0.4905 | -0.2133 | 0.8313 | -0.5039 | 0.6149 | 0.2187 | 0.8271 |
| FJK | 3.0316 | 0.0028 | 3.2649 | 0.0013 | -0.4862 | 0.6274 | -4.4140 | 1.7E-05 | -2.3501 | 0.0198 | 1.2624 | 0.2083 |
| FLMN | 2.9977 | 0.0031 | 3.5144 | 0.0005 | 1.2404 | 0.2163 | -3.4964 | 0.0006 | -0.2640 | 0.7921 | 3.4151 | 0.0008 |
| HA | 5.1090 | 7.7E-07 | 1.9645 | 0.0509 | -0.7565 | 0.4502 | -3.9518 | 0.0001 | -0.9504 | 0.3431 | 1.6241 | 0.1060 |
| HB | 2.2672 | 0.0245 | 6.4053 | 1.1E-09 | 2.5822 | 0.0105 | -0.0327 | 0.9740 | -4.4866 | 1.2E-05 | -2.9536 | 0.0035 |
| HC | 0.4801 | 0.6317 | 0.9125 | 0.3626 | 1.4243 | 0.1559 | -3.1062 | 0.0022 | -0.4762 | 0.6345 | 1.6089 | 0.1092 |
| IA | 6.2658 | 2.3E-09 | 4.2976 | 2.7E-05 | -1.9267 | 0.0555 | -5.0078 | 1.2E-06 | -3.6220 | 0.0004 | 1.5694 | 0.1182 |
| IB | 2.5681 | 0.0110 | 1.9346 | 0.0545 | -0.7753 | 0.4391 | -2.2301 | 0.0269 | 0.8769 | 0.3816 | 2.6841 | 0.0079 |
| LA1 | -2.5897 | 0.0103 | -1.9424 | 0.0535 | 0.2647 | 0.7915 | 1.4648 | 0.1446 | -1.2374 | 0.2174 | -2.4161 | 0.0166 |
| LA2 | -1.6033 | 0.1105 | -4.0248 | 8.1E-05 | -3.6663 | 0.0003 | 1.2158 | 0.2255 | -4.0126 | 8.5E-05 | -5.1906 | 5.2E-07 |
| LC1 | 5.6146 | 6.7E-08 | 2.5837 | 0.0105 | -0.9654 | 0.3355 | -1.2818 | 0.2014 | -1.9520 | 0.0524 | -1.1603 | 0.2473 |
| LC2 | 4.8113 | 3.0E-06 | -1.4977 | 0.1358 | -4.2126 | 3.8E-05 | -2.0762 | 0.0392 | -2.7337 | 0.0068 | -1.0617 | 0.2897 |
| LC3 | 1.2885 | 0.1991 | -0.9223 | 0.3575 | -1.4873 | 0.1385 | -0.2467 | 0.8054 | -1.7775 | 0.0770 | -1.4357 | 0.1527 |
| LD | -0.1527 | 0.8788 | -0.9507 | 0.3429 | -0.7452 | 0.4570 | -1.2615 | 0.2086 | 1.1579 | 0.2483 | 2.2492 | 0.0256 |
| LE | 4.5366 | 1.0E-05 | -1.1285 | 0.2605 | -1.4853 | 0.1391 | 0.1773 | 0.8594 | -0.3088 | 0.7578 | -0.1964 | 0.8445 |
| OA | 4.5050 | 1.1E-05 | 4.0286 | 8.0E-05 | 1.7992 | 0.0735 | -3.9993 | 9.0E-05 | -4.0062 | 8.8E-05 | -2.6524 | 0.0086 |
| OB | 3.7636 | 0.0002 | 2.6659 | 0.0083 | 0.8129 | 0.4173 | -2.2212 | 0.0275 | -5.2444 | 4.0E-07 | -4.2606 | 3.2E-05 |
| OC | 2.1273 | 0.0346 | 5.8066 | 2.5E-08 | 4.7327 | 4.2E-06 | -0.9432 | 0.3467 | -2.0013 | 0.0467 | -2.0914 | 0.0378 |
| PA | 5.9661 | 1.1E-08 | 5.2249 | 4.4E-07 | 2.1177 | 0.0355 | -6.1022 | 5.5E-09 | -5.6969 | 4.4E-08 | -2.9386 | 0.0037 |
| PB | 4.3414 | 2.3E-05 | 2.6533 | 0.0086 | -0.7040 | 0.4823 | -4.5121 | 1.1E-05 | -3.8278 | 0.0002 | -1.0762 | 0.2832 |
| PC | 5.5920 | 7.5E-08 | 3.7371 | 0.0002 | -0.5544 | 0.5800 | -6.0785 | 6.2E-09 | -3.9692 | 0.0001 | 0.4184 | 0.6761 |
| PD | 6.1568 | 4.1E-09 | 3.8438 | 0.0002 | 1.0448 | 0.2974 | -6.0447 | 7.4E-09 | -2.6770 | 0.0081 | -0.4680 | 0.6403 |
| PE | 3.8958 | 0.0001 | 3.8406 | 0.0002 | 1.4562 | 0.1469 | -2.6072 | 0.0098 | -3.8423 | 0.0002 | -2.8097 | 0.0055 |
| PF | 5.3090 | 3.0E-07 | 4.7374 | 4.1E-06 | 1.7993 | 0.0735 | -5.1372 | 6.7E-07 | -3.0539 | 0.0026 | -0.7366 | 0.4623 |
| PG | 3.2540 | 0.0013 | 4.3299 | 2.4E-05 | 2.0086 | 0.0460 | -4.0114 | 8.6E-05 | -2.5956 | 0.0102 | -0.7761 | 0.4386 |
| PH | 3.7802 | 0.0002 | 4.1181 | 5.6E-05 | 1.9825 | 0.0488 | -3.4831 | 0.0006 | -2.1717 | 0.0311 | -0.6683 | 0.5048 |
| TA | 1.9310 | 0.0549 | 2.2640 | 0.0247 | 1.1614 | 0.2469 | -2.0064 | 0.0462 | -0.8429 | 0.4003 | 0.6561 | 0.5125 |
| TB | 2.6127 | 0.0097 | 2.8982 | 0.0042 | 0.5542 | 0.5801 | -0.6124 | 0.5410 | 0.0855 | 0.9319 | 1.4610 | 0.1456 |
| TC | 3.5940 | 0.0004 | -0.1672 | 0.8674 | -3.5372 | 0.0005 | -1.5091 | 0.1329 | -0.8852 | 0.3771 | 0.7254 | 0.4691 |
| TD | 2.5733 | 0.0108 | 0.4557 | 0.6491 | -1.6707 | 0.0964 | -1.1446 | 0.2538 | 0.4238 | 0.6722 | 1.2706 | 0.2054 |
| TE | 4.3883 | 1.9E-05 | 5.4422 | 1.6E-07 | 2.1425 | 0.0334 | -2.2008 | 0.0289 | -2.2992 | 0.0225 | -1.2125 | 0.2268 |
| TF | 4.1409 | 5.1E-05 | 2.7630 | 0.0063 | 0.3084 | 0.7581 | -1.4515 | 0.1482 | -0.6271 | 0.5313 | -0.0162 | 0.9871 |
| TG | 2.4801 | 0.0140 | 1.9012 | 0.0587 | -1.4811 | 0.1402 | -1.8683 | 0.0632 | -0.9836 | 0.3265 | 2.1208 | 0.0352 |

Note. Regional effects from figure 3C. Green to Red cell shading indicates magnitude, green text color indicates significance at $p < .05$ .

A gradient was observed across the cortex, most notably within models of thickness, demonstrating that primary sensory and motor areas declined the most due to baseline white matter health. This contrasts with that of the main effect of age, likely because age is included in this model but is treated as a regressor when interpreting the main effect of baseline diffusion. Therefore, age accounts for much of the variance observed in these association-like cortices. Once age is accounted for, the remaining variance is primarily in these primary sensory and motor areas, specifically the precentral (FA) and agranular frontal (FB), as well as all the regions encompassing the postcentral (PA, PB, PC, and PD). In fact, many of the prefrontal regions are showing weak or non-significant coupling between each aspect of morphometry with baseline white matter.

Gross differences were observed among morphometric features such that volume and thickness loss was more strongly associated with differences in baseline white matter than was surface area. Differences between thickness and volume measures were minimal but were observed in posterior limbic areas (LC1, LC2, and LE) demonstrating stronger relationships with thickness than volume, and pre-frontal areas demonstrating stronger relationships with volume than thickness (FD, FE, FDT). Unique GM/WM coupling is found in the anterior limbic areas LA1 and LA2 such that slight decreases in morphometric measures were found with higher baseline FA. Surface area showed unique relationships with white matter in several ways. Most notably, coupling with surface area was weaker or non-significant compared to volume and thickness across much of the cortex including the prerolandic frontal, parietal, and temporal areas. Additionally, prefrontal regions show declines in surface area with increased baseline FA that mirror regions that were non-significant in models with thickness.

### 3.5 Change in gray matter morphometry associated with the interaction of age and baseline measures of white matter diffusion

Given that morphometric change was found to be heavily influenced by (coupled with) both age and white matter, we examined the interacting effects of both factors to reveal where coupling of gray and white matter was age-dependent (**Figure 3D**; **Table 6**). Interaction effects were evident throughout the prefrontal and temporal cortices such that lower baseline FA was indicative of greater thickness and volume decline across the lifespan. Similarly, higher MD was indicative of greater morphometric rate of decline across the lifespan. These effects were similar in that they were driven by proxies of degraded white matter health from lower baseline FA and higher baseline MD. Additionally, these results follow similar gradients to those observed in the main effects such that the most significant interactions were in regions responsible for higher-order association processing. Moving along the cortex, away from tertiary hubs towards primary regions in the frontal and temporal lobes, resulted in attenuated or non-significant effects. Interestingly, anterior limbic regions demonstrated similar coupling across the lifespan for volume and surface area metrics. In the LA1 and LA2 regions, lower baseline WM values led to greater morphometric declines across the lifespan. These results are intuitive for FA, but unexpected for MD as typically lower MD would indicate healthier baseline WM. In these regions we found no significant interactions as thickness declined at a similar rate across the lifespan regardless of baseline levels of WM.

**Table 6.**
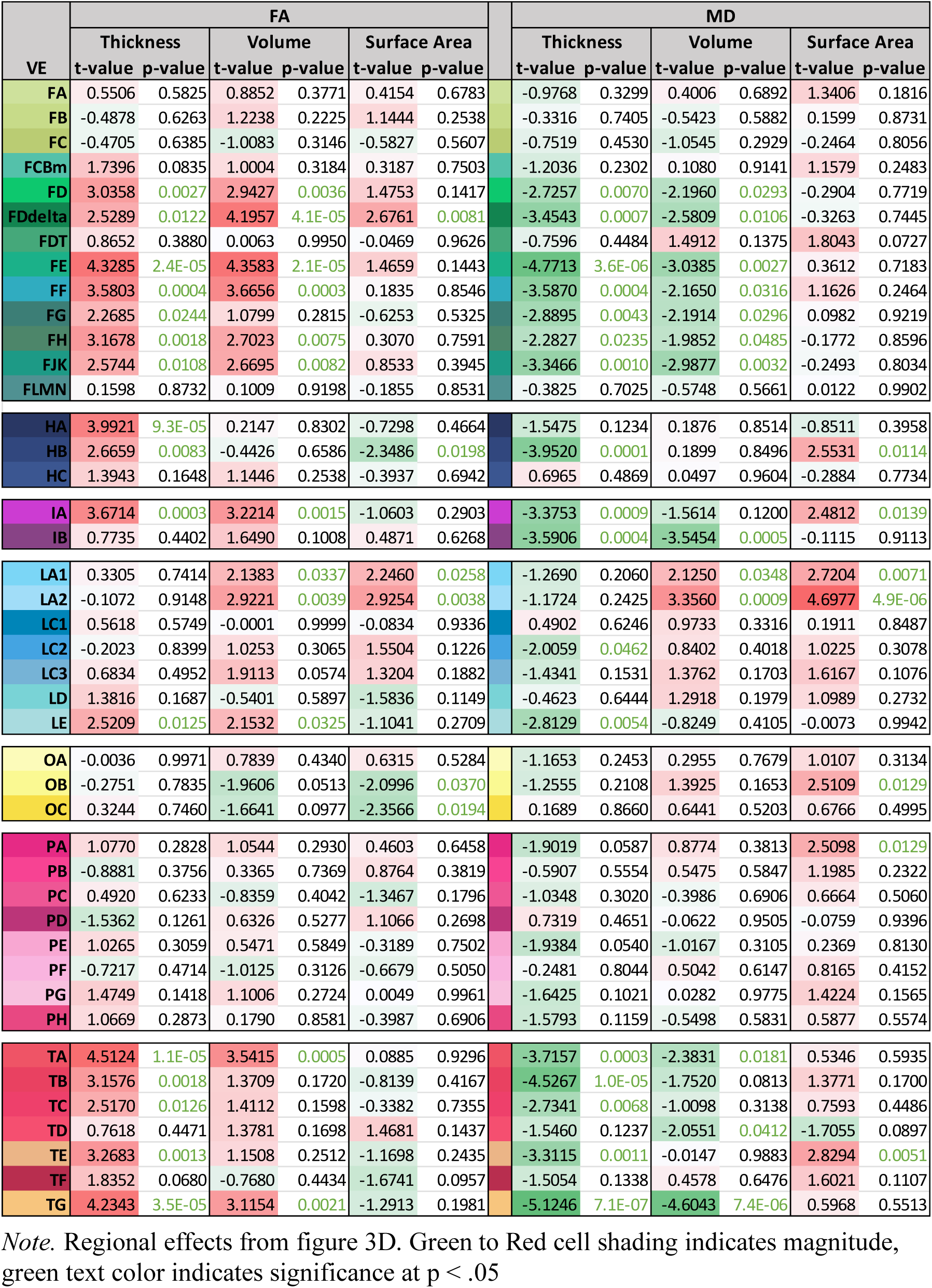
Gray matter change associated with the interaction between age and baseline white matter.

| VE | FA |  |  |  |  |  | MD |  |  |  |  |  |
| --- | --- | --- | --- | --- | --- | --- | --- | --- | --- | --- | --- | --- |
|  | Thickness |  | Volume |  | Surface Area |  | Thickness |  | Volume |  | Surface Area |  |
|  | t-value | p-value | t-value | p-value | t-value | p-value | t-value | p-value | t-value | p-value | t-value | p-value |
| FA | 0.5506 | 0.5825 | 0.8852 | 0.3771 | 0.4154 | 0.6783 | -0.9768 | 0.3299 | 0.4006 | 0.6892 | 1.3406 | 0.1816 |
| FB | -0.4878 | 0.6263 | 1.2238 | 0.2225 | 1.1444 | 0.2538 | -0.3316 | 0.7405 | -0.5423 | 0.5882 | 0.1599 | 0.8731 |
| FC | -0.4705 | 0.6385 | -1.0083 | 0.3146 | -0.5827 | 0.5607 | -0.7519 | 0.4530 | -1.0545 | 0.2929 | -0.2464 | 0.8056 |
| FCBm | 1.7396 | 0.0835 | 1.0004 | 0.3184 | 0.3187 | 0.7503 | -1.2036 | 0.2302 | 0.1080 | 0.9141 | 1.1579 | 0.2483 |
| FD | 3.0358 | 0.0027 | 2.9427 | 0.0036 | 1.4753 | 0.1417 | -2.7257 | 0.0070 | -2.1960 | 0.0293 | -0.2904 | 0.7719 |
| FDdelta | 2.5289 | 0.0122 | 4.1957 | 4.1E-05 | 2.6761 | 0.0081 | -3.4543 | 0.0007 | -2.5809 | 0.0106 | -0.3263 | 0.7445 |
| FDT | 0.8652 | 0.3880 | 0.0063 | 0.9950 | -0.0469 | 0.9626 | -0.7596 | 0.4484 | 1.4912 | 0.1375 | 1.8043 | 0.0727 |
| FE | 4.3285 | 2.4E-05 | 4.3583 | 2.1E-05 | 1.4659 | 0.1443 | -4.7713 | 3.6E-06 | -3.0385 | 0.0027 | 0.3612 | 0.7183 |
| FF | 3.5803 | 0.0004 | 3.6656 | 0.0003 | 0.1835 | 0.8546 | -3.5870 | 0.0004 | -2.1650 | 0.0316 | 1.1626 | 0.2464 |
| FG | 2.2685 | 0.0244 | 1.0799 | 0.2815 | -0.6253 | 0.5325 | -2.8895 | 0.0043 | -2.1914 | 0.0296 | 0.0982 | 0.9219 |
| FH | 3.1678 | 0.0018 | 2.7023 | 0.0075 | 0.3070 | 0.7591 | -2.2827 | 0.0235 | -1.9852 | 0.0485 | -0.1772 | 0.8596 |
| FJK | 2.5744 | 0.0108 | 2.6695 | 0.0082 | 0.8533 | 0.3945 | -3.3466 | 0.0010 | -2.9877 | 0.0032 | -0.2493 | 0.8034 |
| FLMN | 0.1598 | 0.8732 | 0.1009 | 0.9198 | -0.1855 | 0.8531 | -0.3825 | 0.7025 | -0.5748 | 0.5661 | 0.0122 | 0.9902 |
| HA | 3.9921 | 9.3E-05 | 0.2147 | 0.8302 | -0.7298 | 0.4664 | -1.5475 | 0.1234 | 0.1876 | 0.8514 | -0.8511 | 0.3958 |
| HB | 2.6659 | 0.0083 | -0.4426 | 0.6586 | -2.3486 | 0.0198 | -3.9520 | 0.0001 | 0.1899 | 0.8496 | 2.5531 | 0.0114 |
| HC | 1.3943 | 0.1648 | 1.1446 | 0.2538 | -0.3937 | 0.6942 | 0.6965 | 0.4869 | 0.0497 | 0.9604 | -0.2884 | 0.7734 |
| IA | 3.6714 | 0.0003 | 3.2214 | 0.0015 | -1.0603 | 0.2903 | -3.3753 | 0.0009 | -1.5614 | 0.1200 | 2.4812 | 0.0139 |
| IB | 0.7735 | 0.4402 | 1.6490 | 0.1008 | 0.4871 | 0.6268 | -3.5906 | 0.0004 | -3.5454 | 0.0005 | -0.1115 | 0.9113 |
| LA1 | 0.3305 | 0.7414 | 2.1383 | 0.0337 | 2.2460 | 0.0258 | -1.2690 | 0.2060 | 2.1250 | 0.0348 | 2.7204 | 0.0071 |
| LA2 | -0.1072 | 0.9148 | 2.9221 | 0.0039 | 2.9254 | 0.0038 | -1.1724 | 0.2425 | 3.3560 | 0.0009 | 4.6977 | 4.9E-06 |
| LC1 | 0.5618 | 0.5749 | -0.0001 | 0.9999 | -0.0834 | 0.9336 | 0.4902 | 0.6246 | 0.9733 | 0.3316 | 0.1911 | 0.8487 |
| LC2 | -0.2023 | 0.8399 | 1.0253 | 0.3065 | 1.5504 | 0.1226 | -2.0059 | 0.0462 | 0.8402 | 0.4018 | 1.0225 | 0.3078 |
| LC3 | 0.6834 | 0.4952 | 1.9113 | 0.0574 | 1.3204 | 0.1882 | -1.4341 | 0.1531 | 1.3762 | 0.1703 | 1.6167 | 0.1076 |
| LD | 1.3816 | 0.1687 | -0.5401 | 0.5897 | -1.5836 | 0.1149 | -0.4623 | 0.6444 | 1.2918 | 0.1979 | 1.0989 | 0.2732 |
| LE | 2.5209 | 0.0125 | 2.1532 | 0.0325 | -1.1041 | 0.2709 | -2.8129 | 0.0054 | -0.8249 | 0.4105 | -0.0073 | 0.9942 |
| OA | -0.0036 | 0.9971 | 0.7839 | 0.4340 | 0.6315 | 0.5284 | -1.1653 | 0.2453 | 0.2955 | 0.7679 | 1.0107 | 0.3134 |
| OB | -0.2751 | 0.7835 | -1.9606 | 0.0513 | -2.0996 | 0.0370 | -1.2555 | 0.2108 | 1.3925 | 0.1653 | 2.5109 | 0.0129 |
| OC | 0.3244 | 0.7460 | -1.6641 | 0.0977 | -2.3566 | 0.0194 | 0.1689 | 0.8660 | 0.6441 | 0.5203 | 0.6766 | 0.4995 |
| PA | 1.0770 | 0.2828 | 1.0544 | 0.2930 | 0.4603 | 0.6458 | -1.9019 | 0.0587 | 0.8774 | 0.3813 | 2.5098 | 0.0129 |
| PB | -0.8881 | 0.3756 | 0.3365 | 0.7369 | 0.8764 | 0.3819 | -0.5907 | 0.5554 | 0.5475 | 0.5847 | 1.1985 | 0.2322 |
| PC | 0.4920 | 0.6233 | -0.8359 | 0.4042 | -1.3467 | 0.1796 | -1.0348 | 0.3020 | -0.3986 | 0.6906 | 0.6664 | 0.5060 |
| PD | -1.5362 | 0.1261 | 0.6326 | 0.5277 | 1.1066 | 0.2698 | 0.7319 | 0.4651 | -0.0622 | 0.9505 | -0.0759 | 0.9396 |
| PE | 1.0265 | 0.3059 | 0.5471 | 0.5849 | -0.3189 | 0.7502 | -1.9384 | 0.0540 | -1.0167 | 0.3105 | 0.2369 | 0.8130 |
| PF | -0.7217 | 0.4714 | -1.0125 | 0.3126 | -0.6679 | 0.5050 | -0.2481 | 0.8044 | 0.5042 | 0.6147 | 0.8165 | 0.4152 |
| PG | 1.4749 | 0.1418 | 1.1006 | 0.2724 | 0.0049 | 0.9961 | -1.6425 | 0.1021 | 0.0282 | 0.9775 | 1.4224 | 0.1565 |
| PH | 1.0669 | 0.2873 | 0.1790 | 0.8581 | -0.3987 | 0.6906 | -1.5793 | 0.1159 | -0.5498 | 0.5831 | 0.5877 | 0.5574 |
| TA | 4.5124 | 1.1E-05 | 3.5415 | 0.0005 | 0.0885 | 0.9296 | -3.7157 | 0.0003 | -2.3831 | 0.0181 | 0.5346 | 0.5935 |
| TB | 3.1576 | 0.0018 | 1.3709 | 0.1720 | -0.8139 | 0.4167 | -4.5267 | 1.0E-05 | -1.7520 | 0.0813 | 1.3771 | 0.1700 |
| TC | 2.5170 | 0.0126 | 1.4112 | 0.1598 | -0.3382 | 0.7355 | -2.7341 | 0.0068 | -1.0098 | 0.3138 | 0.7593 | 0.4486 |
| TD | 0.7618 | 0.4471 | 1.3781 | 0.1698 | 1.4681 | 0.1437 | -1.5460 | 0.1237 | -2.0551 | 0.0412 | -1.7055 | 0.0897 |
| TE | 3.2683 | 0.0013 | 1.1508 | 0.2512 | -1.1698 | 0.2435 | -3.3115 | 0.0011 | -0.0147 | 0.9883 | 2.8294 | 0.0051 |
| TF | 1.8352 | 0.0680 | -0.7680 | 0.4434 | -1.6741 | 0.0957 | -1.5054 | 0.1338 | 0.4578 | 0.6476 | 1.6021 | 0.1107 |
| TG | 4.2343 | 3.5E-05 | 3.1154 | 0.0021 | -1.2913 | 0.1981 | -5.1246 | 7.1E-07 | -4.6043 | 7.4E-06 | 0.5968 | 0.5513 |

Note. Regional effects from figure 3D. Green to Red cell shading indicates magnitude, green text color indicates significance at $p < .05$

Despite the strength of the morphometric declines in parietal cortices, both across the lifespan and with baseline WM, no parietal regions demonstrated a significant interaction. Closer inspection revealed that parietal regions show consistent declines in morphometry due to baseline diffusion across the lifespan. Other regions showing non-significant effects were typically primary motor, sensory, and visual areas of the post-central, pre-central, and occipital regions.

## 4 DISCUSSION

### 4.1 Overview of results

#### Novelty of study

Building off decades of previous research, the current study substantiates patterns of structural vulnerability of regional gray matter, but also highlights the impact of underlying white matter to neuronal health and the aging process. Previous attempts to concurrently assess white and gray matter structural health in aging included several methods with varying success, and suffered limitations such as use of correlation-based statistical methods or the omission of older adults (Tamnes et al., 2010; Westlye et al., 2010). One group found age-related associations between gray matter thickness and skeletonized white matter with patterns of regional differentiation akin to evolutionary precedence (Kochunov et al., 2011). Advanced statistical modeling has also been utilized, but has failed to link aging to regional differentiation, or isolate tissue types that demonstrate coupled relationships (Groves et al., 2012). Findings from our own cross-sectional research revealed an age-related coupling of gray matter thickness and white matter fractional anisotropy, providing support for retrogenesis-like theories of differential vulnerability, although limited in regional specificity beyond the frontal and parietal lobes (Hoagey et al., 2019).

Here we report 1) within-person declines in cortical thickness and volume across much of the cortex, primarily in frontal, parietal, and temporal lobes; 2) spatial gradients of association including a) regional differentiation as frontal, parietal, and temporal lobes demonstrate the greatest changes, and b) within-region gradients as areas further from sensory and motor hubs demonstrate stronger age-related effects; 3) poorer estimates of white matter health predict greater gray matter morphometric change over time, most prominently in the frontal, parietal, and temporal lobes, suggesting coupled vulnerability in these regions; 4) coupled gray and white matter structural health is increasingly more vulnerable as age increases, specifically in prefrontal and temporal regions, indicating that the biological mechanisms influencing the coupled declines in neuronal health are exacerbated with advanced aging. Additionally, our use of a biologically informed parcellation scheme, the von Economo - Koskinas cyto- and myelo-architectonic informed atlas, enhances interpretation of regional differentiation and sets the stage for future work delineating biological mechanisms of *in vivo* neuroimaging.

### 4.2 Heterochronicity of GM/WM coupling

#### Retrogenesis/SA-axis

Of primary interest are the age-related gradients in structural declines, indicative of differential vulnerabilities in neuronal features. In gray matter, we observed several prominent gradients of association strength such that age-related declines in thickness, volume, and, to a lesser degree, surface area were found to be strongest closer to higher-order cognitive centers. This was most evident, in a tertiary to unimodal fashion, across the frontal lobes from pre-frontal towards pre-rolandic areas, across the parietal lobe from basal and inferior parietal towards the post-central region, and across the temporal lobe from temporal proper towards fusiform, hippocampal, insular, and auditory areas. These findings reiterate previous theories of retrogenesis such that higher-order association cortices develop later in life are the most vulnerable to aging. Conversely, regions closer in proximity to primary motor and sensory hubs demonstrate resilience, exemplifying a sensorimotor-association gradient.

Accounting for the influence of white matter health and assessing the combined impact on changes in gray matter is a key element of our design. As noted, poorer baseline fractional anisotropy was associated with greater declines in gray matter morphometry. Across the lifespan, this was primarily driven by the oldest adults and in frontal and temporal cognitive areas. Overall, this indicates that the biological mechanisms influencing the coupled declines in neuronal health are exacerbated with advanced aging, primarily in association cortices. This may indicate that there is greater age-related loss in the shape or directionality of diffusion as opposed to the magnitude or speed of diffusion. The water flow in the underlying white matter fibers neighboring or adjacent to these gray matter regions, in particular for association cortices, might be experiencing increased dispersion or fanning, such as from axonal thinning or increases in extracellular space, that is exacerbated with aging. While these estimates of white matter structure were taken at baseline, not longitudinally, lower levels of baseline white matter fractional anisotropy in older adults would suggest that this loss of directionality could be underway and predictive of future loss in gray matter morphometry.

Interestingly, parietal regions, which demonstrated strong associations with age and baseline white matter when modeled separately, did not demonstrate age-related interactions to the same degree as frontal and temporal areas. Rather, parietal regions demonstrate consistent linear declines in coupling across the whole adult lifespan. Given that frontal, temporal, and parietal higher-order association areas show coupled declines across both tissue types, it is possible that the mechanisms leading to structural decline occur much earlier and more consistently in parietal regions as opposed to more abrupt declines later in the lifespan in prefrontal and temporal regions. This might suggest differences in the underlying biology of structural declines in the parietal regions compared to frontal and temporal cortices. More research on the specific cytoarchitectural differences or neuronal organization of these regions could clarify these differential patterns further. One highly speculative explanation for these results, given the directionality of our models, is that parietal neurons degrade from cell body to axons (e.g., as in Wallerian degeneration). Given the differential association with diffusion that appears to be informative in determining morphometric declines in frontal and temporal regions, it is possible that frontal and temporal neurons could degrade from cell axons to body (e.g., transneuronal atrophy). As such, baseline white matter health could be predictive of future gray matter morphometric decline in frontal and temporal regions but not parietal. This idea was initially noted in our previous cross-sectional work (Hoagey et al., 2019) and these longitudinal results might provide further evidence to this claim that parietal cortices age differently in their gray-white coupling than other association cortices, although more research is needed with longitudinal data across both modalities to properly tease apart these lead-lag associations. The parietal association cortices are also the regions demonstrating the most protracted development in the brain, and have arisen relatively recently from an evolutionary perspective, which adds credence to the retrogenesis hypothesis of brain aging.

### 4.3 Ubiquitous coupling vs regional differentiation

Directly modeling the relationship between gray and white matter structural health was one of the major goals of the study. We found that there is a coupled relationship between neuronal subparts as cortical regions show change in volume or thickness in association with the baseline diffusion values in adjacent white matter. This was evident across much of the cortex, possibly evidencing ubiquity in the biological connectedness of these tissue proxies as hypothesized. The main effect model accounted for chronological age; therefore, these coupling effects were above and beyond chronological aging. Volumetric differences appeared to be the most pronounced, and this was driven primarily by changes in thickness but not surface area. It is likely that myelin in neighboring white matter leads to differences in thickness of the lamina that does not impact the surface area directly. While this could be due to an inability of the imaging proxies to accurately differentiate between tissue types (Westlye et al., 2009), given that these are acquired in separate imaging sequences, this is better explained as the interplay between neuronal layers and the health of innervating myelin. If the white matter neighboring and innervating the gray matter is thinning, losing axon caliber, or the extracellular space is expanding and taking the place of healthy portions of myelinated axons, this would affect the neighboring gray matter and manifest as a loss of volume, primarily due to the shrinking of laminae and would lead to an overall loss of thickness, without necessarily influencing the cortical surface area. Increased dispersion has been shown to be an indicator of age-related declines in more advanced biophysical multicompartment models. In fact, it was shown that fractional anisotropy shares a large amount of variance with estimates of orientation dispersion, more so than with other estimates of white matter health such as neurite density or myelin water fraction (Billiet et al., 2015). Reduced directionality, or an overall increase in the dispersion of diffusion in these areas at the gray/white interface, could be indicative of initial declines in one of these processes in the neighboring gray matter, leading to a subsequent age-related loss in volume, but specifically in aspects of thickness contributing to overall volume. One exception to this pattern appears to be in the pre-frontal areas as volumetric change seems to be more influenced by decreases in surface area, as there are very few pre-frontal regions showing cortical thinning, beyond age effects. In general, these results support one of our main hypotheses in that the relationship between gray matter and white matter is only somewhat regionally differential, but is better described as ubiquitous, as accounting for the effects of age results in strong coupling between the two tissue compartments across the majority of the cortex. Particularly, these results highlight the coupled relationship between gray matter (specifically volume and thickness) and white matter (specifically fractional anisotropy) structural estimates. Previous cross-sectional findings from our lab have identified a similar pattern of coupled age-related decline of gray matter and white matter diffusion, particularly in the parietal and frontal regions (Hoagey et al., 2019). Differences in our results here are likely due to statistical modeling factors, as chronological age is included in the longitudinal models in the current study, rather than being expressed as a latent component in the partial least squares analysis of the cross-sectional data.

### 4.4 Underlying biology w/Von Economo

#### Regional differences align with myelo/cyto-architecture

The Von Economo – Koskinas atlas, derived from the cellular characteristics and laminar differences among regions, aligns with the regional differentiation in our results matching the original seven-lobe scheme, but also the within-lobe parcellations determined from regional myeloarchitectural and cytoarchitectural features. These sub-divisions map with other recently derived histological and neuroimaging-based features supporting these gradients of age-related change. Regions with a simple laminar structure, primarily heterotypical and agranular homotypic areas, are not as vulnerable to aging processes as homotypic areas with well-defined 6-layer laminar composition. This suggests that the age-related association between gray matter morphometry and neighboring white matter is greater in homotypic association cortices than in heterotypic or agranular regions. The amount of heavily packed granule cells appears to be important compared to areas with less density or the relative absence of granular cells within regions. Dense granular layers are typically those in sensory areas such as visual and somatosensory cortices. Originally termed “koniocortex”, these regions are specialized receptive fields with very dense granular layers optimized for continuous communication with the thalamus and requiring limited flexibility or adaptability over time (Triarhou, 2007). In contrast, the absence of granule cells in layer IV results in larger, long-range connections of the motor and neighboring agranular cortices. These differences in cellular composition directly impact cellular features such as dendritic density and caliber, axon branching, and intracortical myelination. Association cortices with a low density of granule cells allows for complexity in laminar structure and promotes the propagation of larger dendritic arbors to maximize short range connections. The proportion of high caliber projection fibers decreases in regions farther from primary motor and sensory centers (Nieuwenhuys & Broere, 2017). In general, these patterns align with our findings showing a disposition towards worsening structural health across the lifespan in areas comprised of a lower density of granule cells, characteristic of frontal and parietal cortical types in higher-order association cortices. These cortical areas have decreased neuron density due to a high composition of larger neurons, vast dendritic arbors, and modest axon calibers, characteristics typical of 6-layer homotypic cortical types that promote specialized higher-order function and integration across the cortex (Tomer et al., 2022).

Finally, grouping of laminar properties into homotypic and heterotypic categories relates to phylogenetic and ontogenetic cortical patterns that map with our results. Maturation processes can be summarized as a cascade of structural growth with white and gray matter expansion starting with essential visual, motor, and sensory areas developing first, higher-order cognitive centers continuing development throughout childhood, with some areas not fully maturing until early adulthood (Casey et al., 2005). Aging processes mirror those of maturation such that areas that develop latest exhibit greater vulnerability in aging (Raz, 2000; Salat et al., 2004; Scheibel et al., 1975; Yeatman et al., 2014). Laminar complexity aligns with this gradient as the regions with less complex laminar structure, such as agranular and primary areas, complete development first, while more complex areas, such as the von Economo – Koskinas labeled “frontal” and “parietal” cortical types (comprised of prefrontal, temporal, and lateral parietal), develop last. Similarly, the cortical types that are simpler are phylogenetically older while the more elaborate are evolutionarily recent (García-Cabezas et al., 2019), suggesting that the processes leading to evolutionarily newer cortical structure also require longer to develop. Given that these regions also show the most (or earliest) age-related vulnerability, it is likely that this unique laminar composition is responsible for early susceptibility. This “complexity” is often accompanied by high plasticity, an increase in small, tangentially oriented fibers, and the expansion of dendritic arbors within cortical layers. Relatedly, some of the first neurons to migrate from the ventricular germinal zone are radially oriented, followed by the expansion of tangentially oriented fibers and dendritic growth (Marín-Padilla, 1992). While robust intercortical radial and tangential fibers seem to maintain health during aging, an interesting finding in our results is that the earliest signs of aging appear in areas with more tangentially oriented fibers, especially those with extensive dendritic arborization—a pattern noted in previous reports (McNab et al., 2013; Scheibel et al., 1975). Similarly, comparing evolutionary change between a fully developed primate brain to the human neocortex illustrate the expansion of higher-order cognitive regions of the prefrontal, parietal, and lateral temporal cortices (Grydeland et al., 2019).

### 4.5 Limitations

While previous literature details morphometric declines across the lifespan, the current models account for additional factors that add to our understanding of gray matter structural declines. Primarily, morphometric declines are modeled with two time points of longitudinal data while accounting for the baseline differences in white matter health proxies of the neighboring voxels. Given the interconnectedness of these structural proxies, it is important to show that while accounting for variations in white matter health, these morphometric changes remain. Our results highlight our extensive main effects demonstrating morphometric two-wave longitudinal decline, however, there are no notable interactions of inter-wave-interval with either white matter, age, or the combination of white matter and age. This is likely a result of having (thus far) only longitudinal data from two timepoints separated by four years. Change within this short period may not contain enough variance to have differential effects on other variables. Future follow-up analyses with additional timepoints will expand on these limited findings, hopefully bolstering our ability to detect longitudinal change. Additionally, diffusion acquisition sequences and diffusion tensor imaging reconstruction schemes have several pitfalls (Jones & Cercignani, 2010). Analyzing the white matter directly adjacent to cortical gray matter offers a unique perspective of white matter health that might ameliorate some concerns inherent to tract tracing methods, but these voxels are not representative of any specific long range fiber bundles or functional connections between regions and, in fact, might be more related to u-shaped close ranged connections (Catani et al., 2012; Yeterian et al., 2012). Superficial white matter voxels are also more prone to artifacts such as decreased diffusion, directionality, and gyral biases (Reveley et al., 2015) which could influence our results, although we would expect any effect to be consistent and not introduce an age or regional bias. Finally, we do not claim to be measuring any biological or anatomical features directly. Any mention of cytoarchitectural, myeloarchitectural, or cellular features are derived from previous histological or postmortem work, primarily the parcellation and characterization of the von Economo – Koskinas atlas used throughout the study.

## 5 CONCLUSION

Results demonstrate that coupled declines of gray and white matter structural health exhibit differential aging trajectories across the cortex. Gradients of association strength across these proxies of structural vulnerability align with known cortical characteristics and functional hierarchies such that higher-order association-like cortices demonstrate age-related coupled declines in structural health. Overall, these findings highlight the importance of understanding the fine-grained neuronal properties underlying neuroimaging proxies and point toward the importance of integrating cytoarchitecture and myeloarchitectural features in aging research.

## Acknowledgements

This work was funded in part by the National Institutes of Health grant 2R01AG-056535, and foundation grants from AWARE and BvB.

## Data Availability

The data that support the findings of this study are available in the supplementary material of this article.

