## Supplemental Materials for "Coupled aging of cyto- and myeloarchitectonic atlas-informed gray and white matter structural properties"

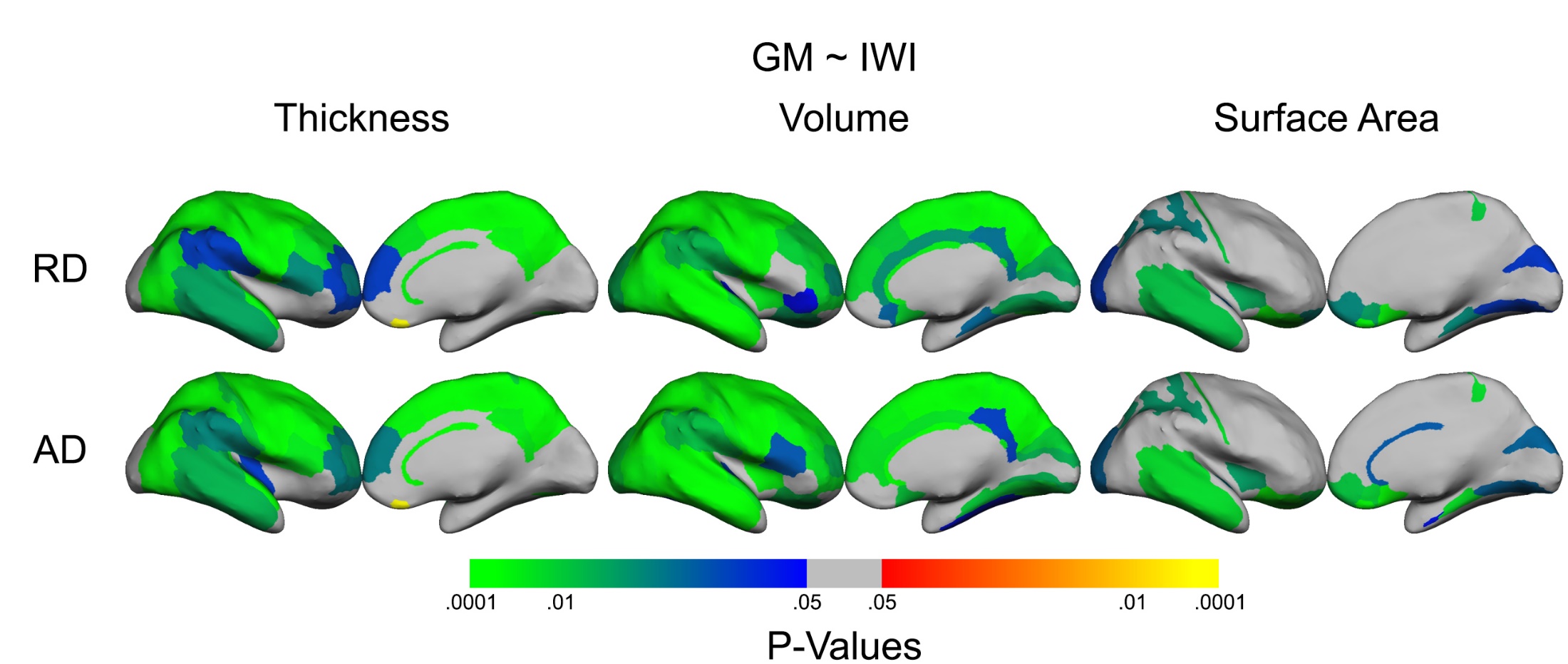
**SUPPLEMENTAL MATERIALS**

Figure S1) Regional main effects of each gray matter morphometric measure across the inter-wave-interval plotted on the inflated von Economo – Koskinas surface

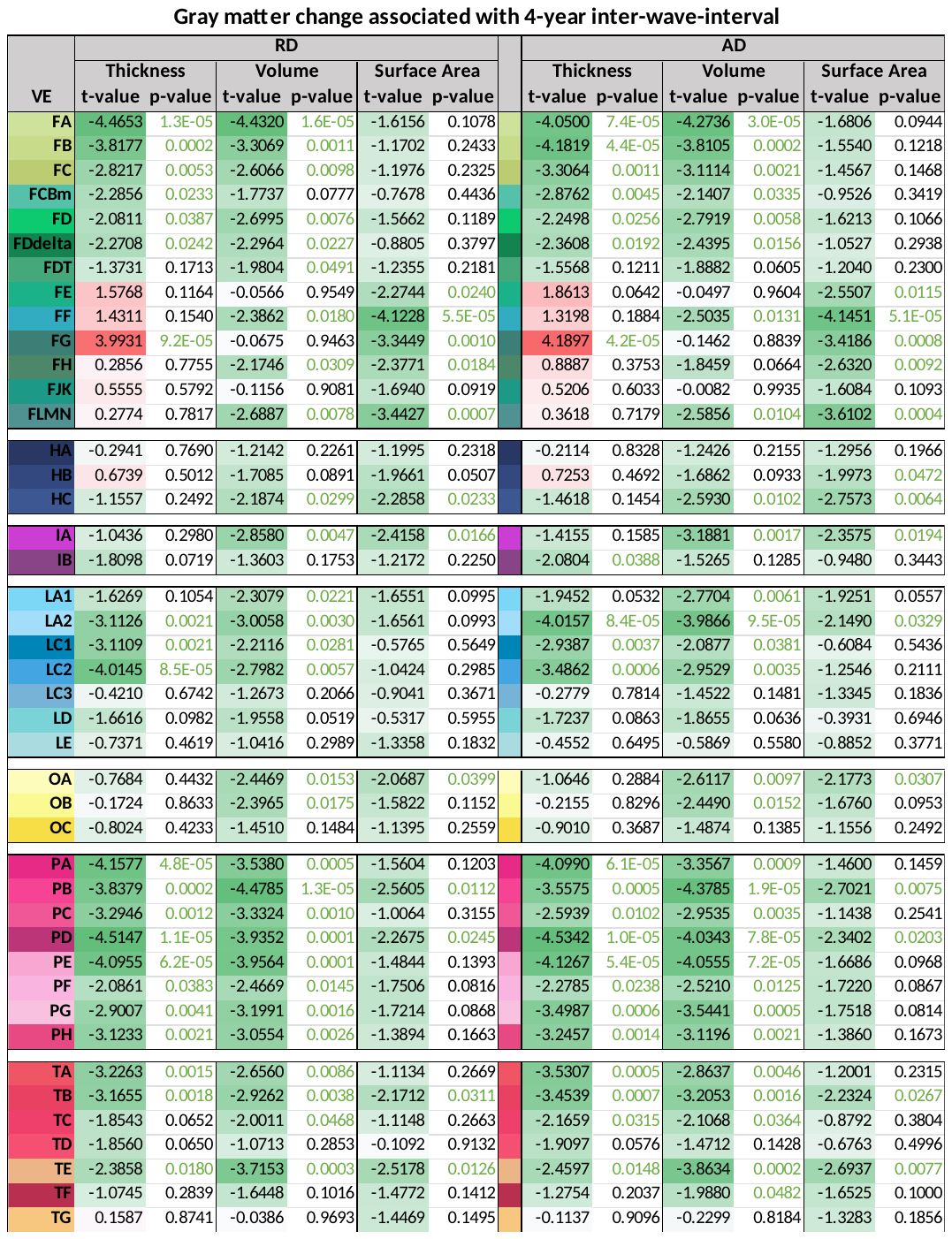

Table S1

Note. Regional linear mixed effects model results from figure S1. Green to red cell shading indicates magnitude, green text color indicates significance at p < .05.

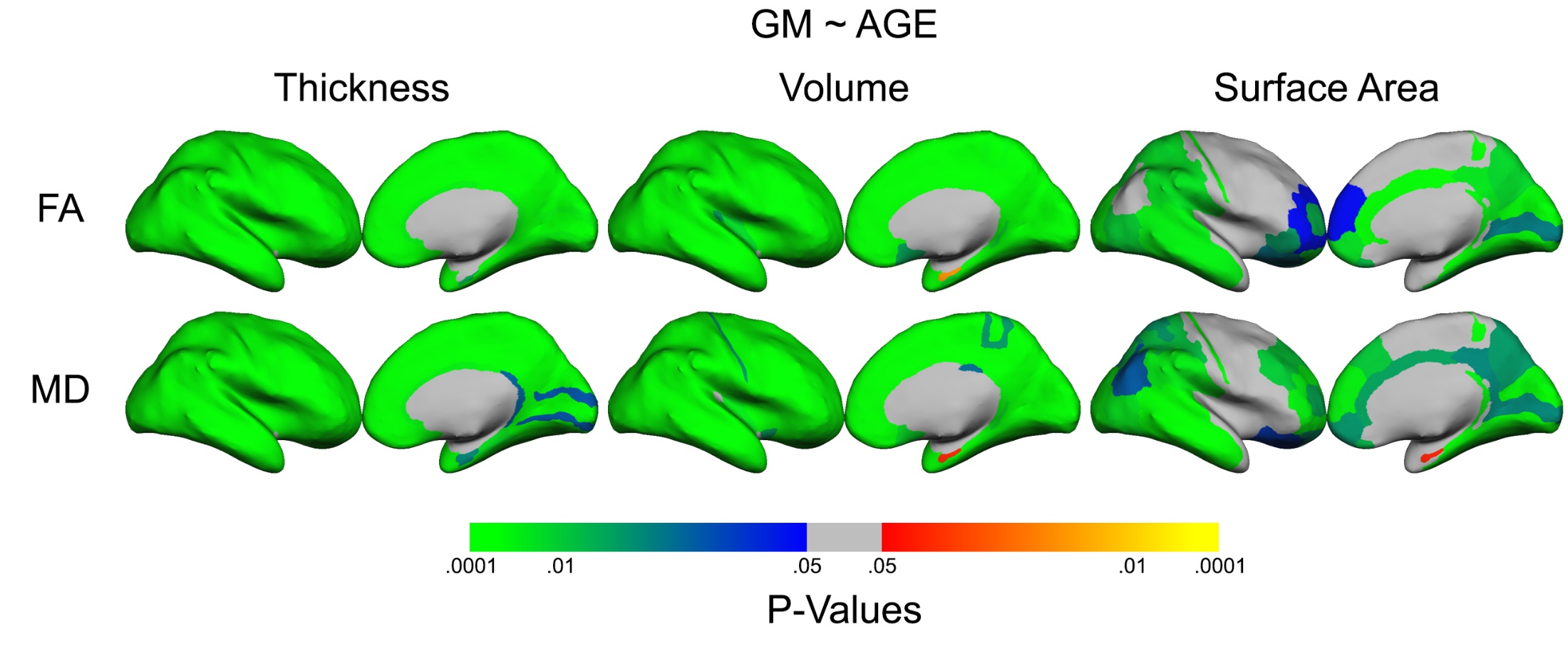

Figure S2) Regional main effects of each gray matter morphometric measure across age plotted on the inflated von Economo – Koskinas surface

Table S2

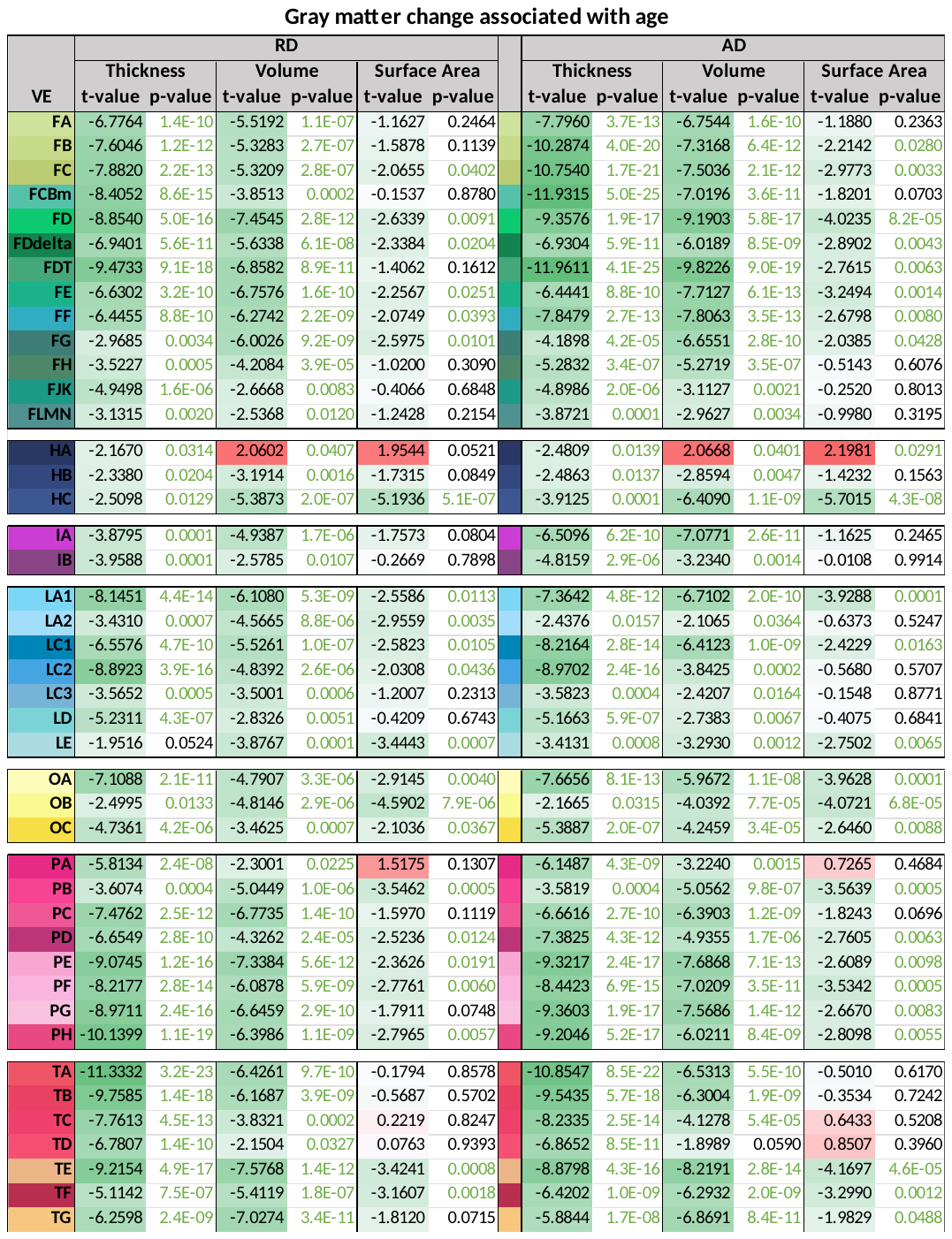

Note. Regional linear mixed effects model results from figure S2. Green to red cell shading indicates magnitude, green text color indicates significance at p < .05.

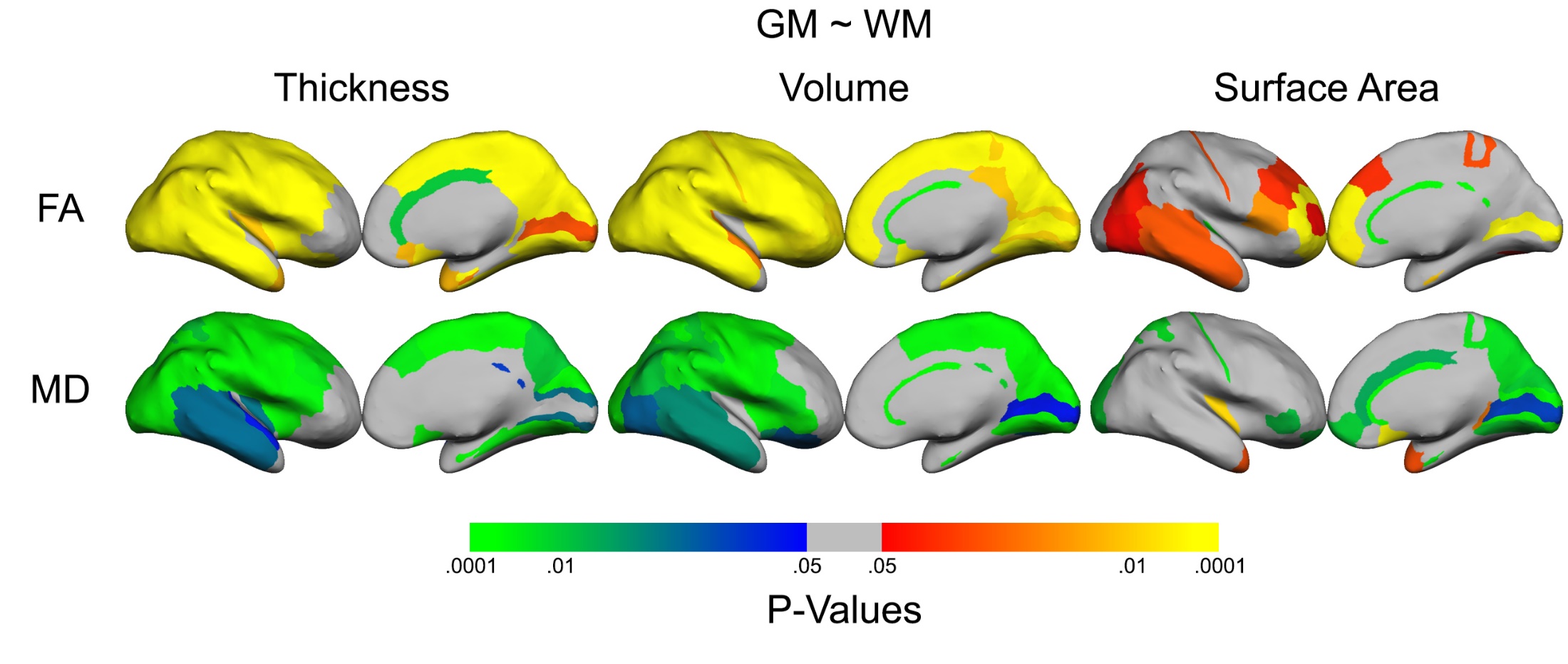

Figure S3) Regional effects of each gray matter morphometric measure across each white matter metric plotted on the inflated von Economo – Koskinas surface

Table S3

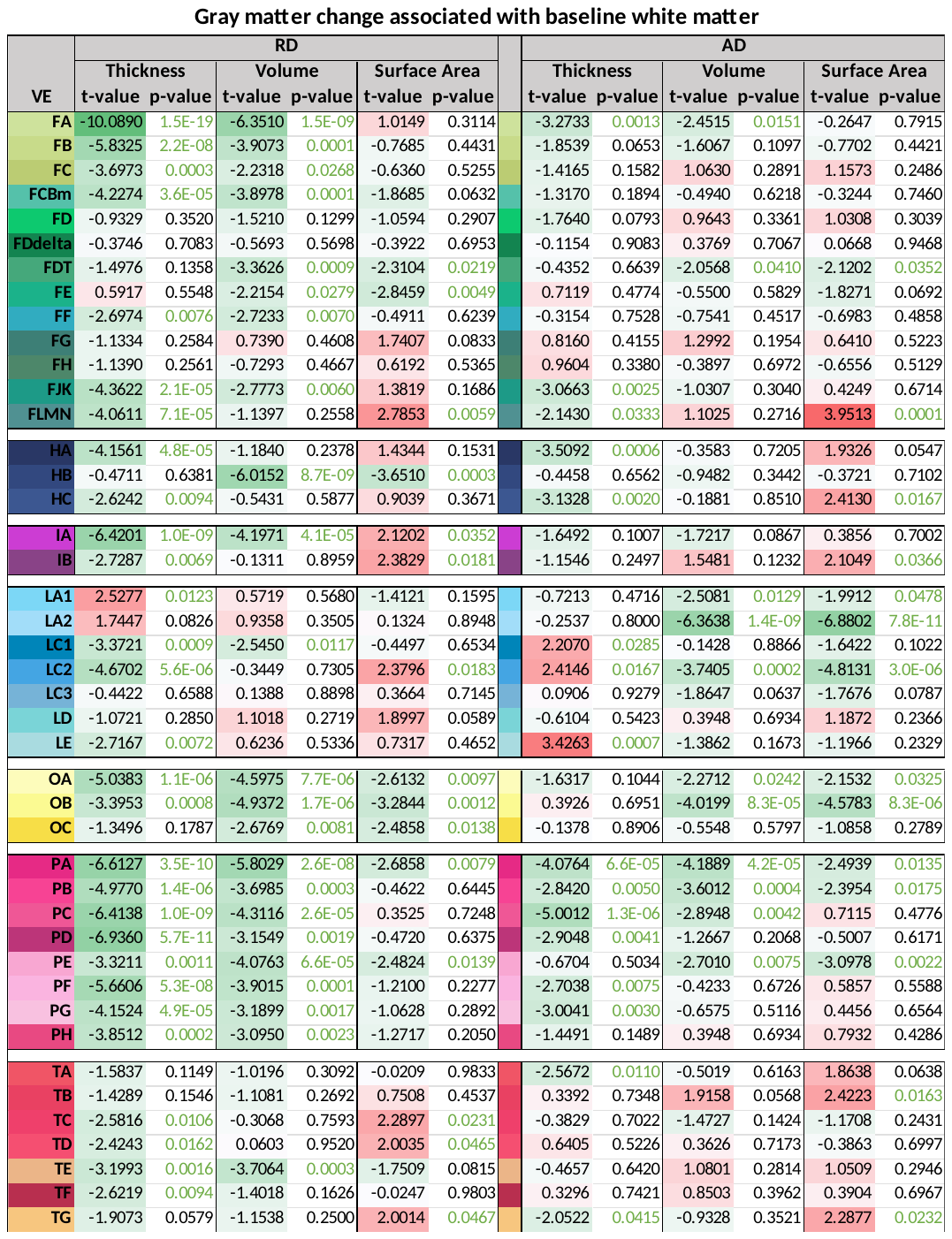

Note. Regional linear mixed effects model results from figure S3. Green to red cell shading indicates magnitude, green text color indicates significance at p < .05.

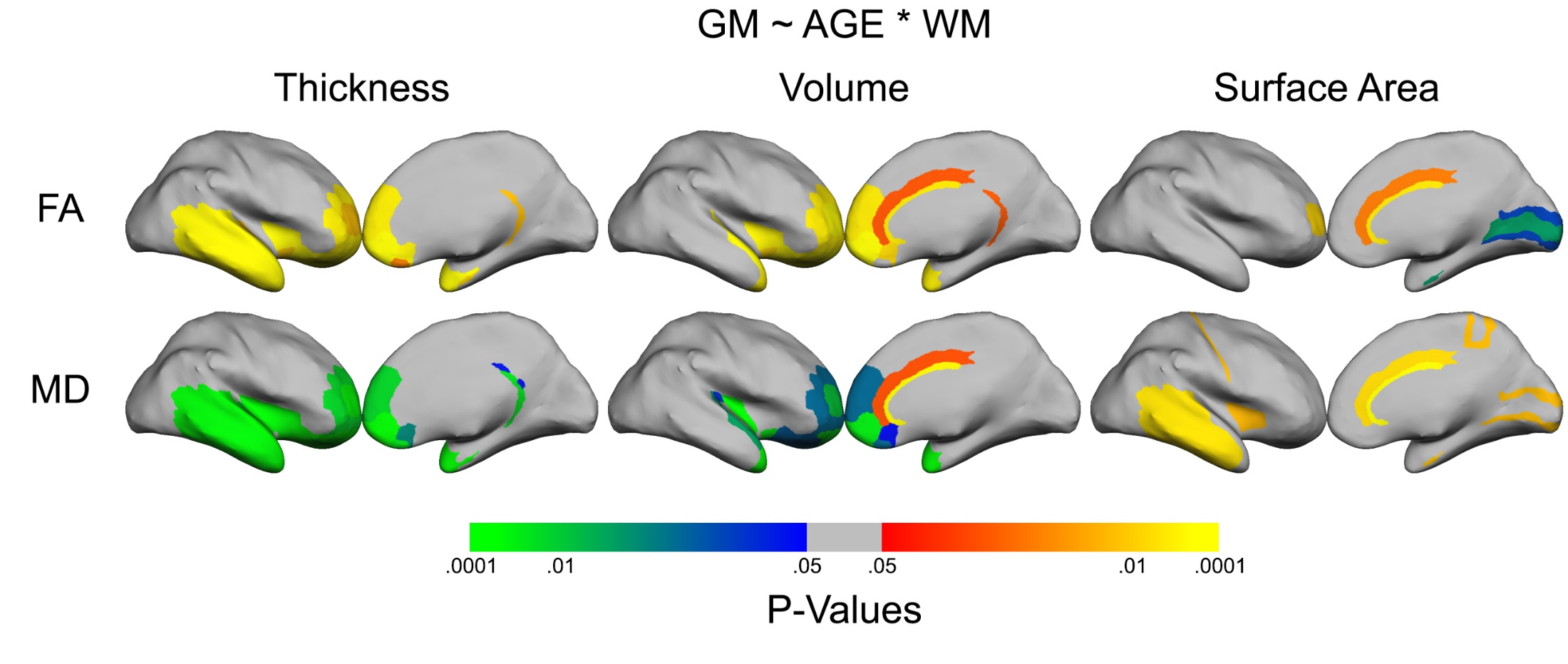

Figure S4) Interaction effects of each gray matter morphometric measure across age and each white matter metric plotted on the inflated von Economo – Koskinas surface

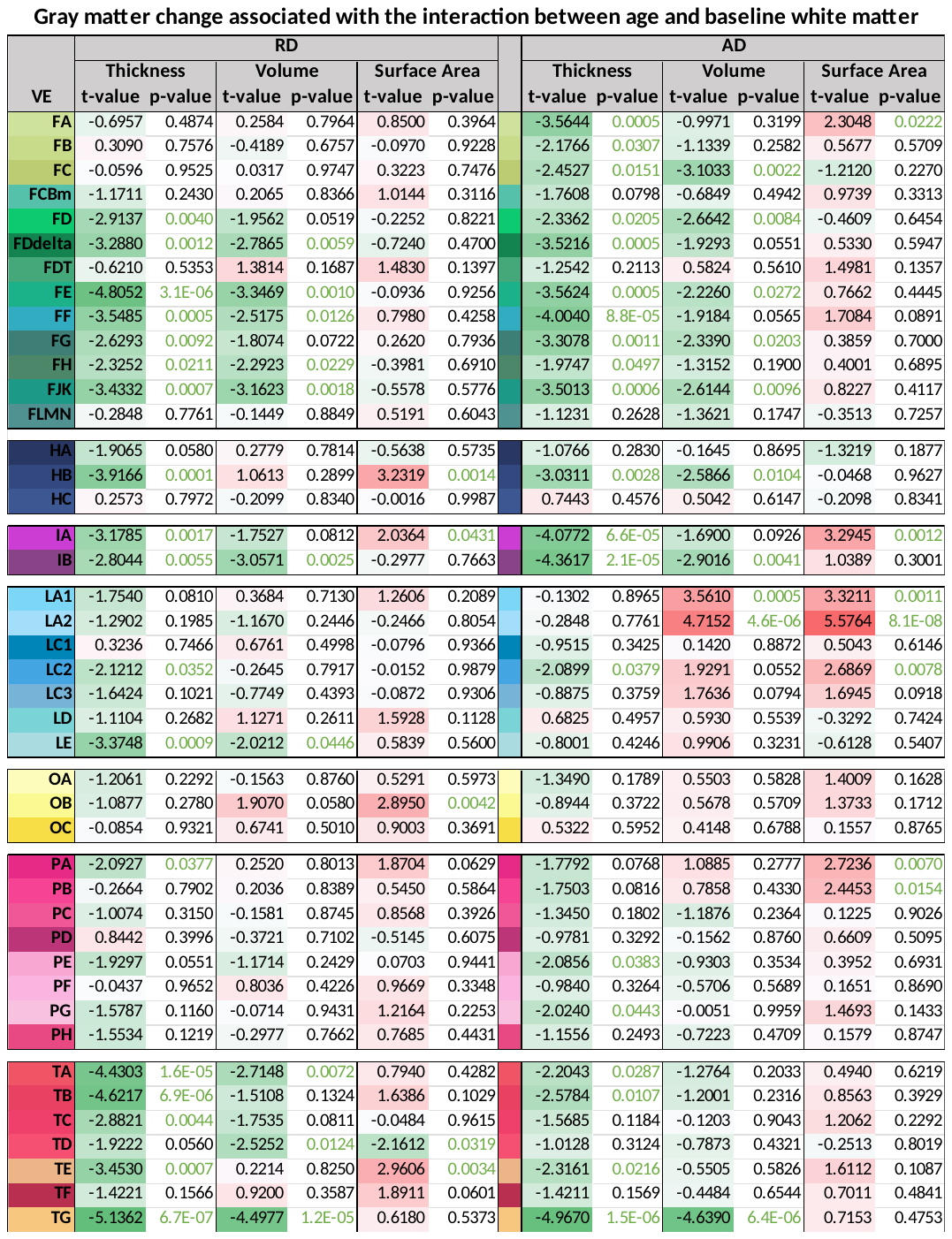

Table S4

Note. Regional linear mixed effects model results from figure S4. Green to red cell shading indicates magnitude, green text color indicates significance at p < .05.

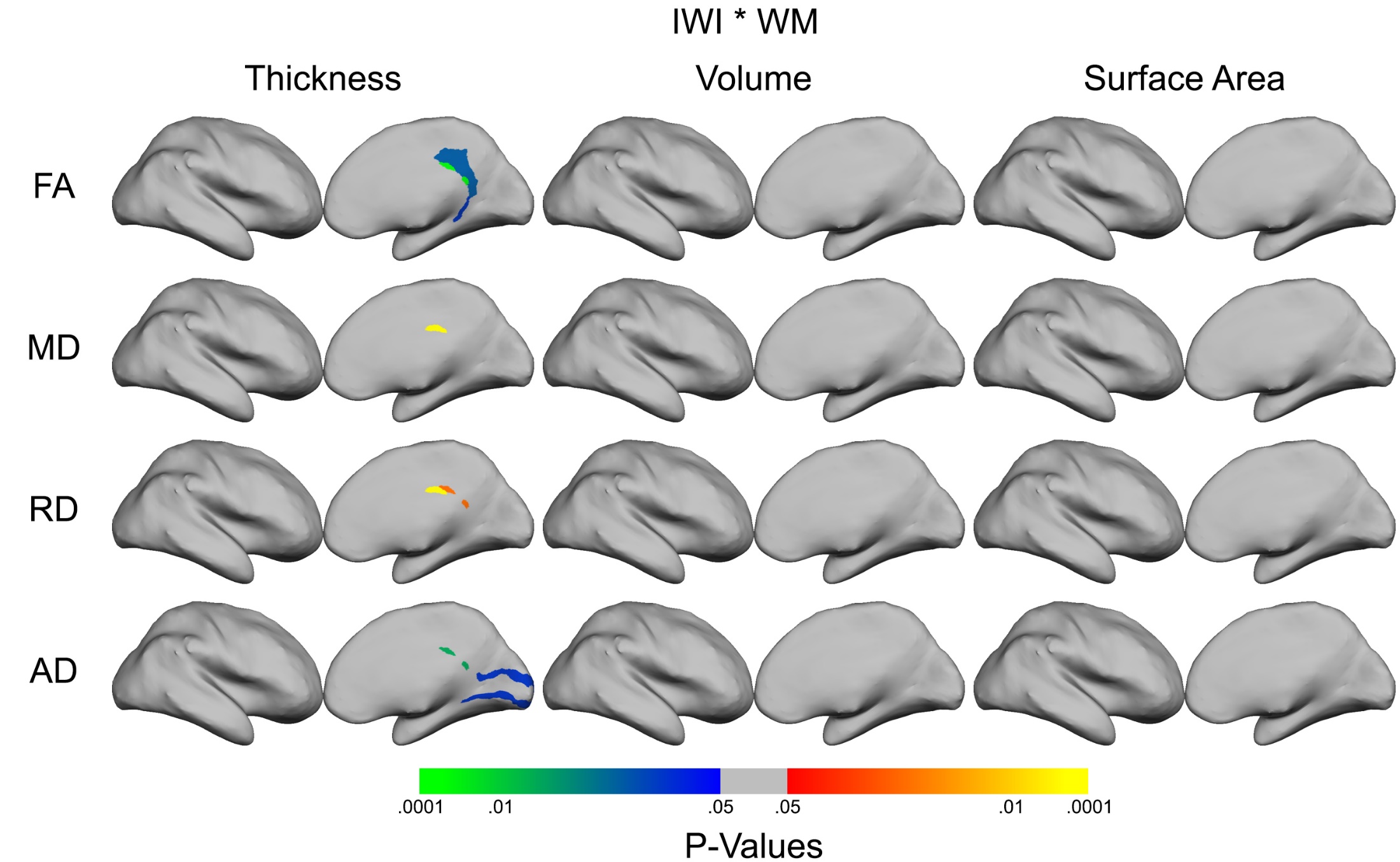

Figure S5) Interaction effects of each gray matter morphometric measure across inter-wave-interval and each white matter metric plotted on the inflated von Economo – Koskinas surface

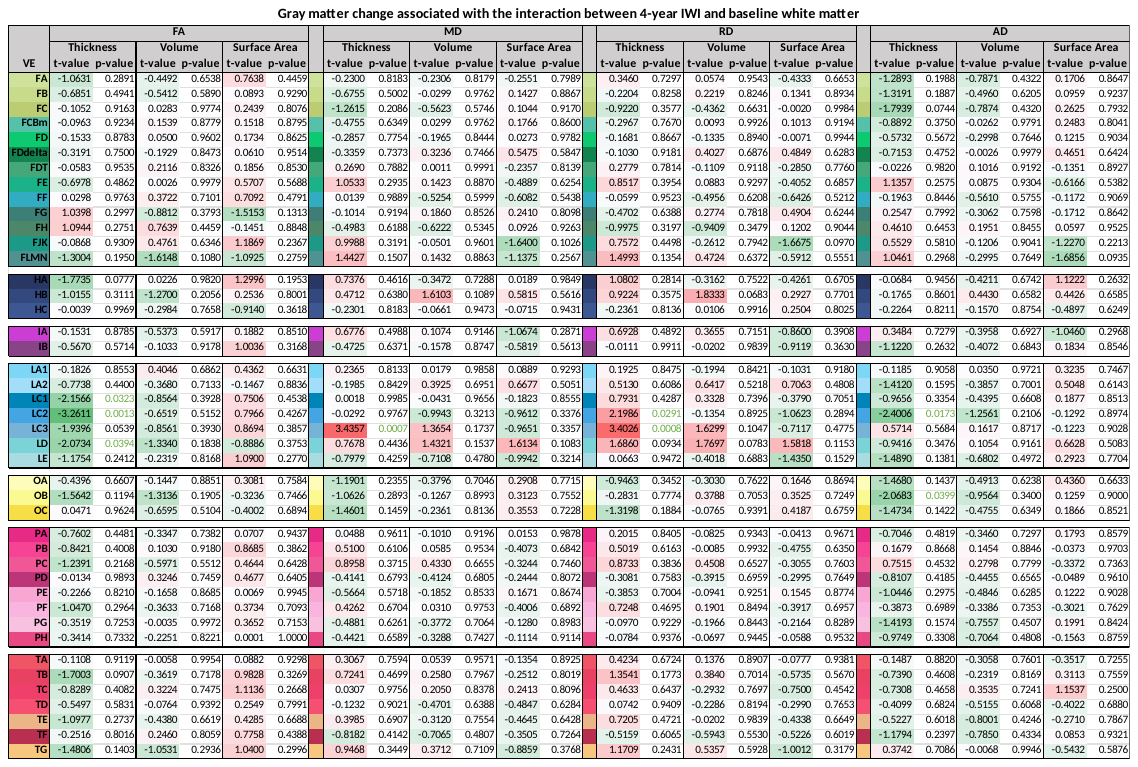

Table S5

Note. Regional linear mixed effects model results from figure S5. Green to red cell shading indicates magnitude, green text color indicates significance at p < .05.

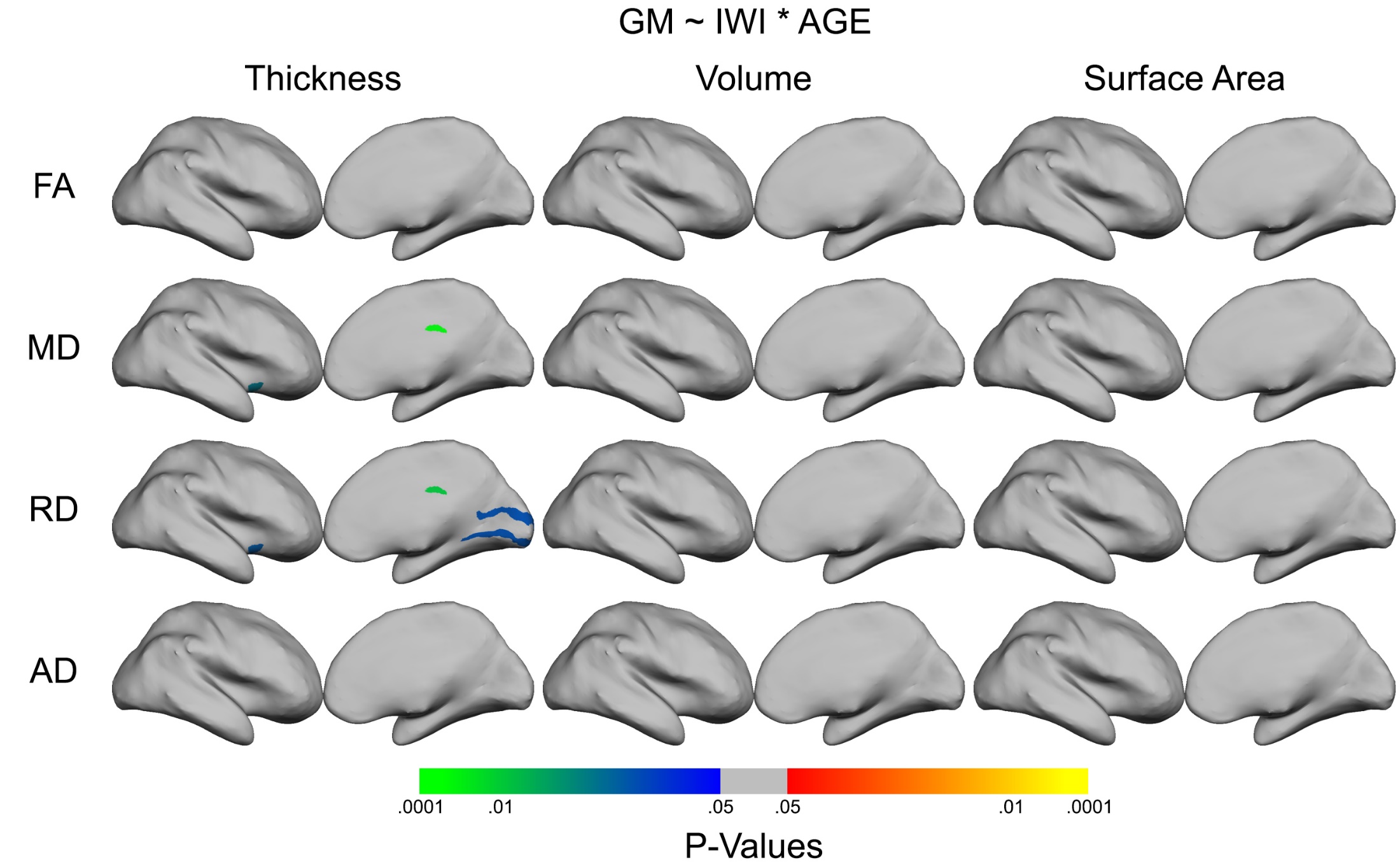

Figure S6) Interaction effects of each gray matter morphometric measure across inter-wave-interval and age plotted on the inflated von Economo – Koskinas surface

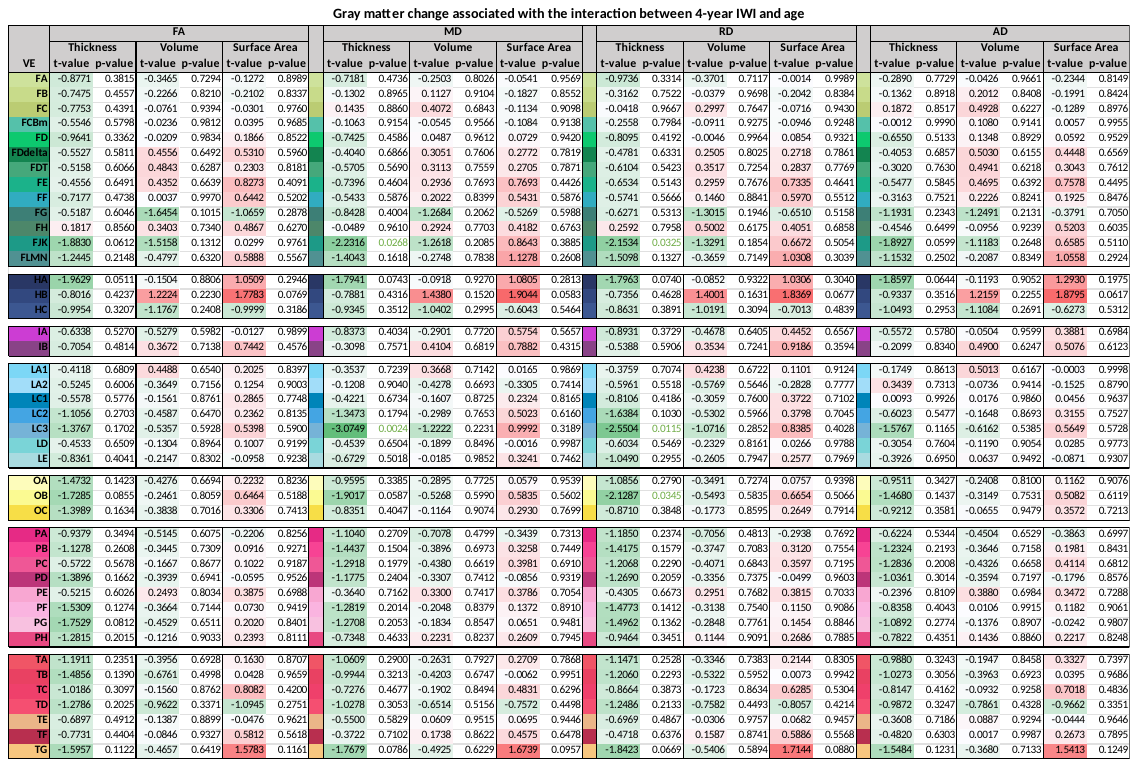

Table S6

Note. Regional linear mixed effects model results from figure S6. Green to red cell shading indicates magnitude, green text color indicates significance at p < .05.

Figure S7) Interaction effects of each gray matter morphometric measures across inter-wave-interval, age, and white matter metric plotted on the inflated von Economo – Koskinas surface

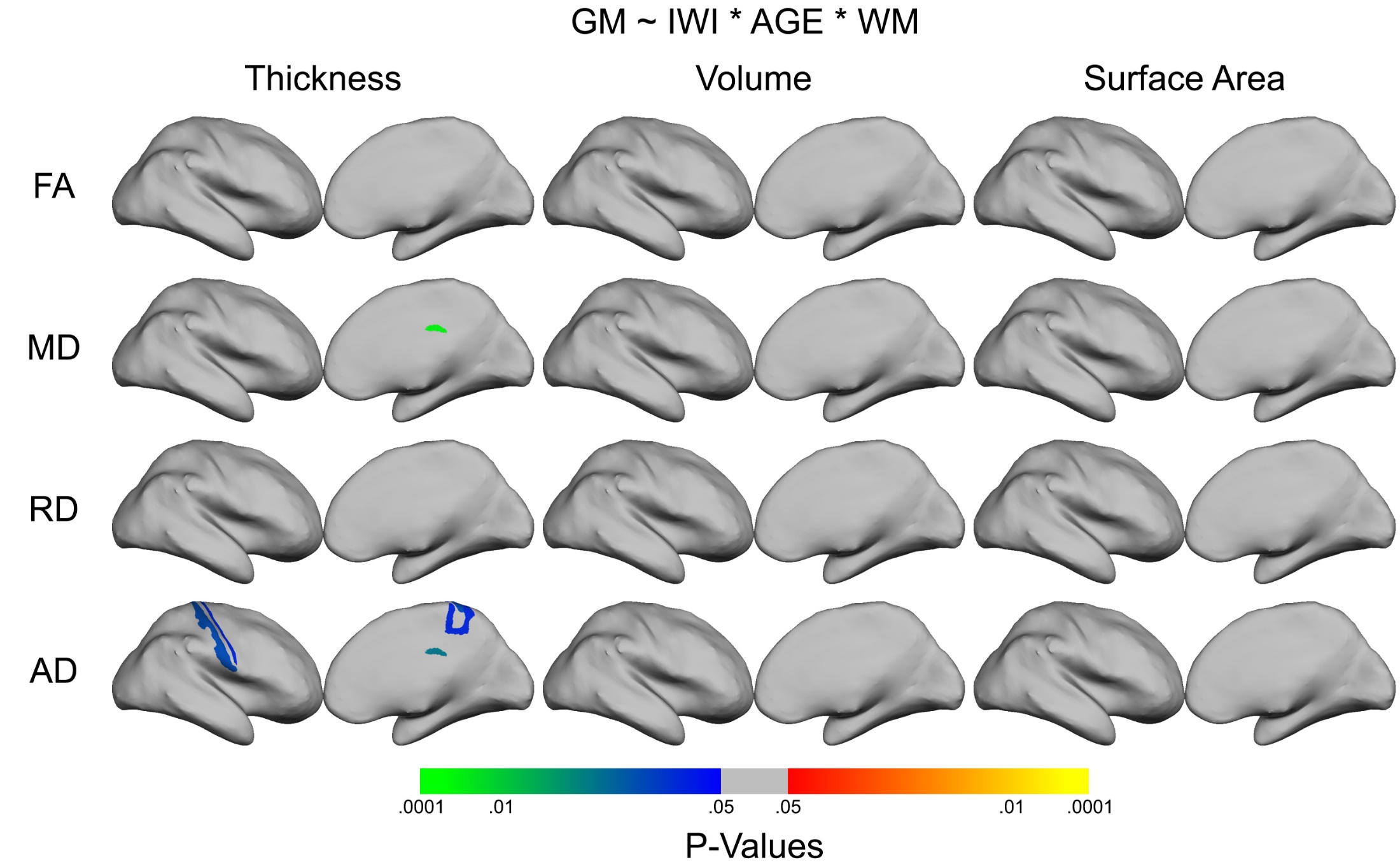

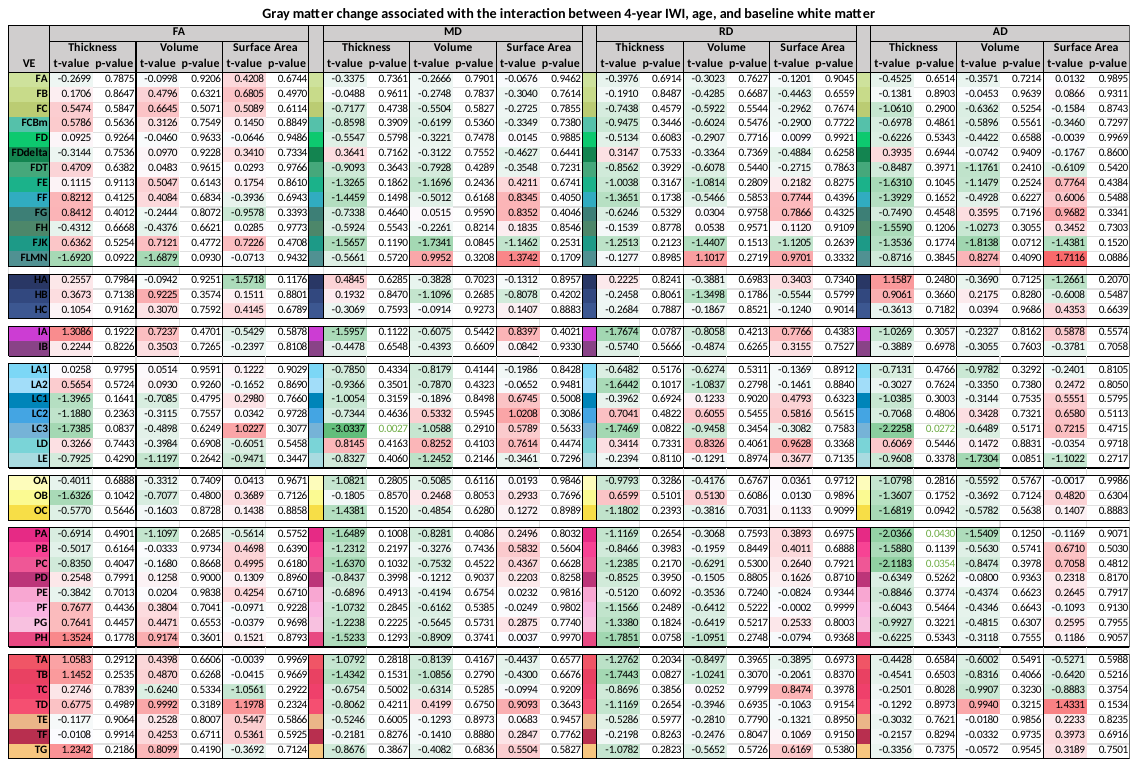

Table S7

Note. Regional linear mixed effects model results from figure S7. Green to red cell shading indicates magnitude, green text color indicates significance at p < .05.
